# Immunomodulation of the Innate Host Response by Mesenchymal-Derived Versican during Influenza A Virus Infection

**DOI:** 10.1101/2025.08.01.668194

**Authors:** Jourdan E. Brune, Mary Y. Chang, Fengying Tang, Cecilia Lopez-Martinez, Stephen R. Reeves, Christina K. Chan, Peter Waldron, David F. Boyd, Sina A. Gharib, Paul G. Thomas, William A. Altemeier, Charles W. Frevert

## Abstract

Viral and bacterial lung infections place a significant burden on public health. Versican, an extracellular matrix (ECM) chondroitin sulfate proteoglycan, coordinates the innate immune response in multiple experimental models. Versican’s potential as an immunomodulatory molecule makes it a promising therapeutic target for controlling the host’s immune response to lung infection. However, versican’s contribution to lung inflammation, injury, and immune cell activity during influenza A virus (IAV) infection represents a critical knowledge gap. To address our central hypothesis that mesenchymal-derived versican is pro-inflammatory and enhances the innate immune response to IAV infection, we generated a tamoxifen-inducible mouse deficient in mesenchymal-derived versican (B6. Col1a2-Cre^ERT+/-^/Vcan^tm1.1Cwf^, Col1a2/Vcan^-/-^). We report that mesenchymal-derived versican plays a critical role in neutrophil, monocyte, and dendritic cell migration into the lungs and airways early in IAV infection. Intriguingly, mesenchymal-derived versican deficiency had the most substantial negative impact on neutrophil emigration into the lungs. We found that neutrophils were less adhesive to the ECM of Col1a2/Vcan^-/-^ mouse lung fibroblasts (mLFs), which had a significant decrease in versican compared to wild-type mLFs. Additionally, Col1a2/Vcan^-/-^ mLFs treated with poly(I:C) *in vitro* have reduced cell-associated hyaluronan. These findings suggest that fibroblast-derived versican is necessary for adhesion to lung fibroblasts by neutrophils as they transit into the lung interstitium and airways from the pulmonary vasculature. Our findings demonstrate that mesenchymal-derived versican is a key regulator of the early host immune responses to IAV.

**NEW & NOTEWORTHY:** We report the novel finding that mesenchymal-derived versican is critical for neutrophil, monocyte, and dendritic cell migration into the lungs and airways early in influenza A virus infection. Additionally, a differentiated neutrophil-cell line is less adherent to versican-deficient fibroblasts, and versican-deficient fibroblasts have significantly reduced cell-associated hyaluronan (HA) content *in vitro*. These findings suggest that mesenchymal-derived versican and cell-associated HA are necessary for the adhesion of neutrophils and monocytes to lung fibroblasts.

## INTRODUCTION

Influenza A virus (IAV) is a major causative pathogen of hospitalization from community-acquired pneumonia and places a significant burden on global health.(1, 2) Globally, an estimated 145,000 to 645,000 annual deaths are attributed to IAV infection.(3, 4) Vulnerable patient populations experience morbidity and mortality associated with seasonal IAV annually, while intermittently, pandemic IAV strains pose a significant health threat to the general population. The current disease burden from seasonal and pandemic influenza A viruses and the growth of vulnerable patient populations underscore the need for continued influenza A research. The extracellular matrix (ECM) is increasingly recognized for its role in maintaining pulmonary tissue homeostasis and coordinating the host immune response to pulmonary diseases.(5–8) Versican, a chondroitin sulfate proteoglycan, is a critical component of the ECM during embryonic development but is present a low levels in healthy, mature lungs.(9) Accumulation of versican and one of its binding partners, hyaluronan (HA), in provisional matrices is a hallmark of many lung diseases.(9–22) Versican is known to interact with multiple receptors found on the surface of immune cells.(8, 20) The accumulation of versican in the lungs and its close association with leukocytes migrating into the pulmonary tissue have been described in C57BL/6J mice infected with a mouse-adapted IAV (A/PR/8/34; H1N1).(23) However, much is still unknown about the immunomodulatory potential of versican and its contextual role in the innate immune response to IAV.

Previous studies utilizing versican-deficient mice have focused on mice globally deficient in versican (Rosa26/Vcan^-/-^), mice deficient in myeloid cell-derived versican (LysM/Vcan^-/-^), and mice deficient in epithelial cell-derived versican (SPC/Vcan^-/-^). These studies demonstrated that Rosa26/Vcan^-/-^ mice treated with polyinosinic:polycytidylic acid (poly(I:C)), a TLR3-agonist, had decreased recovery of inflammatory leukocytes in the bronchoalveolar lavage (BAL) fluid.(24) This contrasts with that of poly(I:C) instilled LysM/Vcan^-/-^ mice, which had increased recovery of inflammatory leukocytes in BAL fluid.(25) Additionally, both Rosa26/Vcan^-/-^ and LysM/Vcan^-/-^ mice had significantly decreased levels of type I interferon and IL-10 in lung tissue and BAL fluid following poly(I:C) exposure compared to wild-type mice.(24, 25) Another study, utilizing SPC/Vcan^-/-^ mice infected with respiratory syncytial virus (RSV), revealed increased migration of neutrophils and monocytes into the BAL fluid and lungs, along with elevated expression of chemokines CCL2, CCL3, and CCL4, suggesting that epithelial cell-derived versican attenuates leukocyte recruitment through chemokine mediated mechanisms.(26) Together, these studies demonstrate the immunomodulatory potential of versican and the importance of the cellular source of versican when considering its role in the host immune response.

It is becoming apparent that mesenchymal cells, which include fibroblasts, smooth muscle cells, and pericytes, play a critical role in the host’s response to pulmonary inflammation and injury.(27–31) Recent studies indicate that fibroblasts create environmental niches that regulate the inflammatory and regenerative responses in the lungs of mice exposed to bleomycin and influenza.(29, 31) Single-cell gene expression studies have expanded our understanding of the heterogeneity of lung fibroblasts and localized them to spatially distinct regions of the lungs.(32–35) Adventitial fibroblasts are localized to the perivascular and peribronchiolar spaces of conducting airways, and alveolar fibroblasts are located in the distal lung near alveolar type II cells.(28, 31) In naïve mice, fibroblasts are the predominant source of versican in the lungs and constitute a significant cell population responsible for versican expression in the lungs under inflammatory conditions.(23, 27, 30) Soucy et al. demonstrated that in response to pneumococcal pneumonia, pulmonary fibroblasts have a distinct transcriptomic signature characterized, in part, by significant induction of versican expression.(30) However, the role of fibroblast-derived versican in other pneumonias remains unknown.

Therefore, this study investigated the role of mesenchymal-derived versican during the host immune response to IAV infection by utilizing a tamoxifen-inducible mouse deficient in mesenchymal-derived versican. To test our hypothesis that mesenchymal-derived versican is pro-inflammatory and enhances the innate immune response to IAV infection, we assessed several metrics of acute lung injury, including leukocyte migration into select pulmonary microenvironments. The underlying mechanisms by which mesenchymal-derived versican alters the innate immune response to IAV infection were explored. We found mesenchymal-derived versican to be critical for neutrophil, monocyte, and dendritic cell migration into the lungs and airways during early IAV infection. Concurrent changes in cytokine and chemokine levels in the BAL fluid were not observed. *In vitro,* neutrophils demonstrated reduced adhesion to versican-deficient mouse lung fibroblasts (mLFs) compared to mLFs from wild-type mice. These findings suggest a pro-inflammatory role for mesenchymal-derived versican within the provisional extracellular matrix during IAV infection.

## MATERIALS and METHODS

### Single-cell RNA Sequencing Dataset Analysis

Using published scRNA-seq data from C57BL/6J mice exposed to PR8 virus for 0, 1, 3, 6, and 21 days post-infection (dpi), we identified mesenchymal cells that expressed Col1a2 and versican.(27) As previously described, the scRNA-seq data was generated using CD45^-^ cells isolated from whole lungs of pathogen-free C57BL/6 mice (Taconic Biosciences) on days 0, 3, and 6 after infection. Whole lungs were collected from four or five mice on each of the three days studied. Mice received an intranasal infection with 2500 EID50 of the mouse-adapted influenza A/Puerto Rico/8/1934 (PR8) virus in 30 µl of 1 x PBS. Single-cell transcriptomic libraries were prepared using the 5′ Gene Expression Kit (V2, 10X Genomics), and sequencing was conducted on the Illumina NovaSeq to produce approximately 500 million reads per sample. Sequencing results were processed using Cell Ranger (v.3.0.2, 10X Genomics) using the mm10 reference altered to include the PR8 influenza genome. After filtering out low quality cells, count data was normalized, and dimensionality reduction and clustering were performed using the R package ‘Seurat’(v.3.0.0.900). Cell clusters were annotated using known markers from the literature. The dataset was later filtered to include only the 0, 3, and 6 time points. (27) For the reanalysis, we generated a subset including those clusters that showed *Col1a1* or *Acta2* expression and repeated the dimensionality reduction and clustering. After reprocessing, expression levels of mesenchymal subpopulation markers and genes of interest were assessed.

### Animals

Vcan^tml.1Cwf^ (Vcan^fl/fl^) mice were bred with mice carrying a transgene for a tamoxifen-inducible Cre recombinase under the control of the Col1a2 promoter (B6.Cg-Tg(Col1a2-Cre/ERT,-ALPP)7Cpd/J (Strain: 029567; Jackson Laboratories) in order to generate B6.Col1a2-Cre^ERT+/-^/Vcan^tm1.1Cwf^ (Col1a2/Vcan^-/-^) mice and B6.Col1a2-Cre^ERT-/-^/Vcan^tm1.1Cwf^ wild-type (WT) littermates.(24) The B6J.B6N(129S4)-Vcan^tm1.1Cwf^/Mmucd mouse strain has been deposited and is available at the Mutant Mouse Resource and Research Centers (MMRRC).

Its Research Resource Identifier is RRID:MMRRC_068231-UCD and Stock Number is 068231-UCD. In Col1a2/Vcan^-/-^ mice, Cre recombinase-mediated deletion of Vcan exon 4 initiates a frameshift, resulting in a STOP codon near the beginning of exon 5. The resulting mice are deficient in mesenchymal-derived versican. All mice were housed under standard conditions in Allentown individually ventilated cages in a specific pathogen-free animal facility. Room lighting was on a 10:14 hr dark:light cycle, and mice had free access to food and water.

Col1a2/Vcan^-/-^ mice and their littermate wild-type controls (WT) were weaned at three weeks of age and fed tamoxifen citrate chow (Inotiv, 400mg/kg, TD.130859, West Lafayette, IN). After being provided tamoxifen citrate chow for four weeks, mice were fed standard mouse chow (LabDiet, PicoLab® Mouse Diet 20 – 5053, St. Louis, MO). At eight-to eleven weeks of age, male Col1a2/Vcan^-/-^ mice and their littermate wild-type controls were infected with a mouse-adapted influenza A virus. All procedures were performed as part of a scientific protocol approved by the University of Washington Institutional Animal Care and Use Committee (IACUC).

### Induction of IAV Pneumonia

Mouse-adapted influenza A/Puerto Rico/8/34 (A/PR/8/34; H1N1) was grown in allantoic fluid of research-grade-specific pathogen-free embryonic chicken eggs (Charles River Avian Vaccine Services, Norwich, CT), and a hemagglutination assay was performed to determine the viral titer.(36) Male mice were infected with 20 plaque-forming units (PFU) in 50 µL PBS by oropharyngeal aspiration under isoflurane anesthesia.(37) This dose caused severe influenza pneumonia, as previously described.(23, 38) Control mice received instillation of PBS alone, and mice were sacrificed at 3, 6, and 9 days post-infection (dpi) by exsanguination under isoflurane anesthesia.

### Quantitative Real-time Reverse-transcription PCR

Mice were sacrificed at 3, 6, and 9 (dpi) with influenza virus A/PR/8/34. Under aseptic conditions, the left lung lobe was removed from the thoracic cavity, used for mRNA isolation, and prepared as previously described.(23) Briefly, lung tissue was placed in 5ml of RNAlater (Invitrogen) at 4C overnight. Lungs were homogenized, and RNA was extracted using RNAeasy Mini Kit with on-column DNase digestion (Qiagen, Valencia, CA) according to the manufacturer’s instructions. cDNA was reverse-transcribed using random primers with the High Capacity cDNA Reverse Transcription Kit (Applied Biosystems, Foster City, CA). Quantitative real-time reverse-transcription polymerase chain reaction (PCR) was performed on an ABI Prism 7900HT Fast Real-Time PCR System (Applied Biosystems) using PrimeTime Gene Expression Master Mix (Integrated DNA Technologies, Coralville, IA). Gene-specific TaqMan primer-probe mixes were used for quantitative real-time PCR of versican (Mm01283063_m1) and TATA-box protein (Mm01277042_m1) mRNA (ThermoFisher Scientific, Grand Island, NY). Quantitative real-time PCR was performed to amplify the matrix protein (*M1*) gene of the PR8 flu virus. The following primers and probes were used: forward: CAGCACTACAGCTAAGGCTATG; reverse: CTCATCGCTTGCACCATTTG, probe /56-FAM/CCTCTGCTG/ZEN/CTTG

CTCACTCGATC/3IABkFQ with 20ng of sample cDNA per reaction. DNA from purified plasmid generated a standard curve for copy number determination. Samples and log10 dilutions of standards were run in triplicate on an ABI 7900 real-time PCR machine, and the viral copy number was determined from the standard curve of Ct values.

### Tissue Harvest and *In Vivo* Antibody Labeling for Spectral Flow Cytometry

Mice were sacrificed at 3 and 6 dpi with influenza virus A/PR/8/34. Before euthanasia, mice were anesthetized with isoflurane in an induction chamber. Alexa Fluor 488-conjugated anti-CD45.2 antibody, clone 104 (BioLegend, San Diego, CA, USA), diluted 1:40 (v:v) with PBS, in a total volume of 200µL (2.5µg/per mouse), was injected retro-orbitally to label the intravascular (IV) leukocytes (IVCD45^+^). After 3 minutes, while keeping the mouse anesthetized with isoflurane delivered by nose cone, the left renal artery was transected, and the mouse was euthanized by exsanguination to reduce the chance that the IVCD45 conjugate would bind non-specifically to airway or interstitial lung leukocytes. In separate experiments investigating circulating leukocytes, mice were euthanized by exsanguination via intracardiac blood draw into an EDTA-coated syringe and did not receive IVCD45 antibody-conjugate injections.

After euthanasia, the trachea was cannulated with an 18-gauge angiocath via tracheostomy, and the airways were immediately lavaged with 1 mL PBS three times. The first 1 mL lavage was separated from the second and third lavages for chemokine and cytokine analysis with multiplex ELISA. The thoracic cavity was opened via midline sternotomy, and the lungs were perfused by injecting 5 mL of PBS into the right ventricle. Lung tissue was removed by blunt dissection from the primary bronchi and placed on ice.

### Preparation of Single-Cell Suspensions for Spectral Flow Cytometry

Cells were prepared for spectral flow cytometry as previously described.(39) Briefly, the first 1 mL of lavage was collected and combined with protease inhibitor according to the manufacturer’s instructions (Pierce™ Protease Inhibitor Moni Tablets, EDTA-Free, A32955, Thermo Fisher Scientific Waltham, MA, USA). These samples were stored at-80°C until chemokine and cytokine analysis. Cells from all three BAL fractions were combined, and RBC lysis buffer (Thermo Fisher Scientific Waltham, MA, USA) was applied according to manufacturer instructions. After RBC lysis, BAL cells were pelleted and suspended in Flow Cytometry Straning (FCS) buffer (Thermo Fisher Scientific Waltham, MA, USA).

Stock solutions of Liberase TM and recombinant DNase I were prepared and stored as previously described.(39) The lungs were minced using a razor blade and then incubated for 45 minutes at 37°C in 2 mL RPMI 1640 without phenol red or FBS (Thermo Fisher Scientific Waltham, MA, USA) containing 0.26U/mL Liberase TM (Sigma-Aldrich, St. Louis, MO, USA) and 10U/mL recombinant DNase I (Sigma-Aldrich, St. Louis, MO, USA). After digestion, samples were pipetted up and down and then filtered through 70µm cell strainers (VWR, Brisbane, CA, USA) with RPMI/10% FBS. Lung cells were pelleted, and RBC lysis buffer was applied (Thermo Fisher Scientific Waltham, MA, USA) according to manufacturer instructions. After RBC lysis, lung cells were pelleted and suspended in an FCS buffer.

Whole blood samples were centrifuged at 2,000x*g* for 15 minutes at 4C. After centrifugation, plasma was combined with protease inhibitor according to the manufacturer’s instructions (Pierce™ Protease Inhibitor Moni Tablets, EDTA-Free, A32955, Thermo Fisher Scientific Waltham, MA, USA) and stored at-80°C until chemokine and cytokine analysis. Blood cells were then combined with 10 mL of Gibco™ ACK Lysing Buffer at room temperature for 4 minutes. Cells were pelleted by centrifugation and combined with 12 mL of cold PBS. Finally, cells were pelleted by centrifugation and suspended in FCS buffer.

### *In Vitro* Antibody Staining for Spectral Flow Cytometry

The spectral flow cytometry panel and staining procedure used have been previously reported elsewhere.(39) Briefly, all antibodies are commercially available, and unstained, single-stain, and fluorescence-minus-one controls were used to define gating for positive and negative populations. Antibody manufacturer, clone numbers, and titers are available in **Table 2**.

**Table 1.** Gating strategy.

| Tissue type | Antigen markers |
| --- | --- |
| Airways | IVCD45<sup>-</sup>CD45<sup>+</sup> collected from bronchoalveolar lavage fluid |
| Lung | IVCD45<sup>-</sup>CD45<sup>+</sup> collected from lung homogenate |
| Marginated vasculature | IVCD45<sup>+</sup>CD45<sup>+</sup> collected from lung homogenate |
| Circulating vasculature | CD45<sup>+</sup> collected from whole blood via cardiac puncture |
| Mediastinal lymph nodes | IVCD45<sup>-</sup>CD45<sup>+</sup> collected from lymph node |
| Cell type | Antigen markers |
| Neutrophils | Ly6G<sup>+</sup> CD24<sup>+</sup> Ly6C<sup>lo/hi</sup> |
| Lymphoid cells | CD64<sup>-</sup> F4/80<sup>-</sup> |
| B cells | CD64<sup>-</sup> F4/80<sup>-</sup> CD19<sup>+</sup> MHCII<sup>+</sup> |
| T cells | CD64<sup>-</sup> F4/80<sup>-</sup> CD3e<sup>+</sup> |
| Cytotoxic T cells | CD64<sup>-</sup> F4/80<sup>-</sup> CD3e<sup>+</sup> CD8<sup>+</sup> |
| T helper cells | CD64<sup>-</sup> F4/80<sup>-</sup> CD3e<sup>+</sup> CD4<sup>+</sup> |
| γδ T cells | CD64<sup>-</sup> F4/80<sup>-</sup> CD3e<sup>+</sup> CD4<sup>-</sup> CD8<sup>-</sup> γδTCR<sup>+</sup> |
| NK cells | CD64<sup>-</sup> F4/80<sup>-</sup> CD3e<sup>-</sup> CD19<sup>-</sup> NKp46<sup>+</sup> CD49b<sup>+</sup> |
| Non-B Non-T Non-NK cells | CD64<sup>-</sup> F4/80<sup>-</sup> CD3e<sup>-</sup> CD19<sup>-</sup> NKp46<sup>-</sup> CD49b<sup>-</sup> |
| Non-lymphoid cells | CD64<sup>+</sup> or F4/80<sup>+</sup> |
| Macrophages | CD64<sup>hi</sup> F4/80<sup>hi</sup> |
| Resident alveolar macrophages<br>(Vehicle instilled mice) | CD64<sup>hi</sup> F4/80<sup>hi</sup> CD11b<sup>-</sup> CD11c<sup>+</sup> SigF<sup>hi</sup> Ly6C<sup>mid</sup> MHCII<sup>-</sup> |
| Resident alveolar macrophages<br>(Influenza infected mice) | CD64<sup>hi</sup> F4/80<sup>hi</sup> CD11b<sup>mid/hi</sup> CD11c<sup>+</sup> SigF<sup>hi</sup> Ly6C<sup>mid</sup> MHCII<sup>-</sup> |
| Recruited macrophages | CD64<sup>hi</sup> F4/80<sup>hi</sup> CD11b<sup>+</sup> CD11c<sup>lo/hi</sup> SigF<sup>lo</sup> Ly6C<sup>hi</sup> MHCII<sup>+</sup> |
| Mono-macrophages | CD64<sup>hi</sup> F4/80<sup>hi</sup> CD11b<sup>+</sup> CD11c<sup>lo/hi</sup> SigF<sup>lo</sup> Ly6C<sup>hi</sup> MHCII<sup>-</sup> |
| Dendritic cells | CD64<sup>lo/int</sup> or F4/80<sup>lo/int</sup> CD11c<sup>+</sup> MHCII<sup>+</sup> |
| CD11b<sup>+</sup> dendritic cells | CD64<sup>lo/int</sup> or F4/80<sup>lo/int</sup> CD11c<sup>+</sup> MHCII<sup>+</sup> CD11b<sup>+</sup> |
| CD103<sup>+</sup> dendritic cells | CD64<sup>lo/int</sup> or F4/80<sup>lo/int</sup> CD11c<sup>+</sup> MHCII<sup>+</sup> CD103<sup>+</sup> |
| Monocytes | CD64<sup>lo/int</sup> or F4/80<sup>lo/int</sup> CD11b<sup>+</sup> CD11c<sup>-</sup> MHCII<sup>-</sup> SigF<sup>-</sup> Ly6C<sup>lo/hi</sup> SSC<sup>lo</sup> |
| Eosinophils | CD64<sup>lo/int</sup> or F4/80<sup>lo/int</sup> CD11c<sup>-</sup> MHCII<sup>-</sup> SigF<sup>+</sup> SSC<sup>hi</sup> |

**Table 2.** Fluorescent Reagents for Cell Staining.

| **Specificity** | **Fluorochrome** | **Clone** | **Vendor** | **Catalog #** | **Titer (ng/test)** |
| --- | --- | --- | --- | --- | --- |
| MHCII-IA/IE | BUV395 | 2G9 | BD Biosciences | 743876 | 8 |
| Viability | Live Dead Blue | - | ThermoFisher | L34962 | 20 μl of 1:320 stock |
| CD103 | BUV737 | M290 | BD Biosciences | 741739 | 125 |
| CD11c | BV421 | N418 | BioLegend | 117330 | 500 |
| Ly6G | Pacific Blue | 1A8 | BioLegend | 127612 | 500 |
| NKp46 | BV605 | 29A1.4 | BD Biosciences | 564069 | 500 |
| Ly6C | BV650 | HK1.4 | BioLegend | 128049 | 31 |
| CD64 | BV711 | X54-5/7.1 | BioLegend | 139311 | 500 |
| Siglec F | BV750 | E50-2440 | BD Biosciences | 747316 | 31 |
| CD4 | BV785 | GK1.5 | BioLegend | 100453 | 63 |
| CD45.2 | Alexa Fluor 488 | 104 | BioLegend | 109816 | 2.5 μg/mouse IV |
| CD45 | PerCP | 30-F11 | BioLegend | 103130 | 125 |
| $\gamma\delta$TCR | PerCP-eFluor710 | GL-3 | ThermoFisher | 46-5711-82 | 63 |
| CD8 | PE | 53-6.7 | BioLegend | 100708 | 63 |
| CD3e | PE-Dazzle 594 | 145-2C11 | BioLegend | 100347 | 500 |
| CD19 | PE-Cy5.5 | 1D3 | ThermoFisher | 35-0193-80 | 500 |
| CD24 | PE-Cy7 | M1/69 | BioLegend | 101822 | 16 |
| F4/80 | Alexa Fluor 647 | BM8 | BioLegend | 123122 | 250 |
| CD11b | APC-R700 | M1/70 | BD Biosciences | 564985 | 16 |
| CD49b | APC-eFluor780 | DX5 | ThermoFisher | 47-5971-82 | 1000 |

Antibodies were diluted in Brilliant Stain Buffer (BD Biosciences San Jose, CA, USA) to achieve the final titers indicated in **Table 2**, allowing for a total volume of 100µL per test. The antibody master mix excluded the antibody conjugate used for *in vivo* staining, Alexa Fluor 488 anti-CD45.2 (IVCD45). For full-stained samples, *in vitro* antibody staining was performed on 1 x 10^6^ cells from the BAL and lung cells of mice that received *in vivo* antibody labeling of IVCD45^+^. BAL and lung cells from a mouse that did not receive IVCD45^+^ labeling were used for the preparation of unstained controls, single-stain viability control, and single-stain cell controls as previously described.(39) Single-stained controls for compatible fluorophore-antibody conjugates were prepared using UltraComp eBeads^TM^ (Thermo Fisher Scientific Waltham, MA, USA), following the manufacturer’s instructions as previously described.(39) *In vitro* antibody staining was performed on single-cell suspensions from whole blood from mice that did not receive *in vivo* antibody labeling of IVCD45^+^ cells. Cells were washed in PBS before incubating in LIVE/DEAD Fixable Blue Dead Cell Stain (Thermo Fisher Scientific Waltham, MA, USA) for 15 minutes. This incubation must be performed in a protein-free solution, without FCS, as protein can impact the efficiency of LIVE/DEAD Blue staining. Next, cells were washed with FCS and incubated with Fc Block TruStain FcX PLUS (anti-mouse CD16/32) (BioLegend, San Diego, CA, USA) diluted 1:100 (v:v) with FCS buffer in a total volume of 50µL per test for 5 minutes. Next, cells were incubated with fluorophore-conjugated antibodies for 30 minutes.

Cells were washed with FCS and then incubated with Cytofix (BD Biosciences San Jose, CA, USA) for 15 minutes. Cells were washed with FCS before being transferred to 5 ml tubes and stored at 4°C in the dark overnight. Flow cytometry was performed the following day.

### Spectral Flow Cytometry

Spectral cytometry was performed using the Aurora 5 laser cytometer (Cytek Biosciences, Fremont, CA, USA) as previously described.(39) Raw data were converted to unmixed data with SpectroFlow software v3.0 using an ordinary least squares algorithm to deconvolute individual fluorophore signatures within a fully stained sample. Identification of multiple autofluorescence (AF) signatures was necessary to unmix specific tissues accurately.(40) AF signatures from unstained BAL, lung, and whole blood tissues were identified and saved as discrete fluorochrome tags in the SpectroFlo library. For all unmixing, the “AF as a tag” feature in the SpectroFlo software was utilized, in addition to incorporating distinct AF spectral signatures into the panel when appropriate. The gating strategy used for these studies has been previously reported.(39) It supports the identification of neutrophils, macrophages, dendritic cells, monocytes, eosinophils, lymphocytes, and natural killer cells. All cytometry data was analyzed using SpectroFlo® (Cytek Biosciences, CA) flow cytometry software.

### Measurement of Total Protein and Inflammatory Mediators in BAL Fluid

Total protein and selected cytokines and chemokines were measured in BAL fluid collected from mice instilled with PBS (0 dpi) and IAV at days 3, 6, and 9 dpi. The total protein concentration of BAL fluid was measured using the Pierce® BCA Protein Assay Kit (Cat. 23225, Thermo Fisher Scientific, Waltham, MA) following the manufacturer’s instructions. Cytokines and chemokine analysis were performed using a Milliplex ZMAP Mouse Cytokine/Chemokine magnetic bead panel (EMD Millipore Corporation, Billeric, MA) with specificity for IL-4, IL-6, IL-10, IL-13, CCL2, CXCL1, CXCL2, TNFα, and VEGF. **CXCL1 and CXCL2 ELISA**

Paired BAL, plasma, and whole lung homogenate samples were collected from mice 3 dpi with IAV to evaluate CXCL1 and CXCL2 chemokine gradients. BAL and plasma samples were collected as described previously in the methods for tissue harvest for spectral flow cytometry and stored at-80°C until analysis. After intravascular perfusion, the left lung was collected by blunt dissection to generate whole lung homogenate. The left lung was placed in 1 mL PBS containing protease inhibitor according to the manufacturer’s instructions (Pierce™ Protease Inhibitor Moni Tablets, EDTA-Free, A32955, Thermo Fisher Scientific Waltham, MA, USA). Lung tissue was homogenized using an Omni Bead Rupter 24 (Omni International, Kennesaw, GA) and 2.8mm Ceramic Beads (Omni International, Kennesaw, GA). After homogenization, samples were aliquoted and stored at-80°C. The DuoSet Elisa Mouse CXCL1/KC and Mouse CXCL2/MIP-2 kits (R&D Systems, Inc. Minneapolis, MN) were used according to the manufacturer’s instructions to evaluate CXCL1 and CXCL2 levels in paired samples.

### Acute Lung Injury Score

Histological assessment of lung inflammation and injury was completed using formalin-fixed paraffin-embedded tissue consisting of the right lung lobes stained with hematoxylin and eosin (H&E).(41) The lung injury score was performed using a modified semiquantitative scoring system as previously described.(23) The analysis was performed by a comparative pathologist (C.W.F) blinded to the mouse genotype and treatment group.

### Stromal Cell Culture

As previously described, mouse lung fibroblasts were isolated from whole lung explants of manipulation-naïve mice.(23, 42) Briefly, lungs were aseptically removed from the thorax, minced, and digested for 60 minutes at 37°C in 2.5 mL DMEM containing 1000 U/mL of Liberase TL and 1000U/mL of DNase I. Digests were filtered through 100µm cell strainer with DMEM containing 10% FBS. The red blood cells were lysed by suspending the cells in sterile water for 30 seconds. Fibroblasts were then plated with DMEM containing 20% FBS, 2 mM L-glutamine, 100 IU/ml penicillin, and 100 µg/ml streptomycin and incubated at 37°C with 5% CO_2_. The cell culture medium was changed every 3–4 days, and fibroblasts were passaged once they were confluent. Fibroblasts were maintained in DMEM 10% FBS, 2 mM L-glutamine, 100 IU/ml penicillin, and 100 µg/ml streptomycin.

For all experiments, fibroblasts were used between passages 2 and 4. For versican western blotting, mouse lung fibroblasts (mLFs) were grown to 80% confluency in 60mm cell culture in 10% FBS DMEM. Cells were serum starved for 24 hours in DMEM without FBS before being incubated with either DMEM 2% FBS with 10 µg/mL of poly(I:C) or fresh DMEM 2% FBS without poly(I:C) for 24 hours. For hyaluronan ELISA and neutrophil adhesion assays, mLFs were seeded in 96-well plates at 3.7 x 10^4^/cm^2^ (1.2 x 10^4^ cells/well) in DMEM 10% FBS and allowed to adhere and grow for 24 hours. Then the medium was changed to DMEM without FBS, and the cells were serum starved for 24 hours. Finally, cells were incubated with either fresh DMEM 2% FBS with 10 µg/mL of poly(I:C) or fresh DMEM 2% FBS without poly(I:C) for 24 hours.

### Versican Western Blot

Versican quantification by western blotting was assessed using a modification of reported methods.(43) Briefly, whole lung tissue and mLF cell preparations were concentrated and purified by ion-exchange chromatography on diethylaminoethyl (DEAE) Sephacel (Sigma-Aldrich, St Louis, MO) in 8M Urea buffer (8M Urea, 2mM EDTA, 50mM Tris base, 0.25M NaCl, 0.5% TX-100, pH 7.5). The columns were washed with 8M Urea buffer and eluted with 8M urea buffer containing 2M NaCl. Samples were ethanol precipitated and digested with 2.0U/mL chondroitinase ABC lyase at 37°C for 3 hours. Samples were loaded under reducing conditions and ran on 4-12% gradient polyacrylamide-SDS gels with 3% polyacrylamide stacking gels overnight at 20V. Samples were then transferred to nitrocellulose and blocked with 10% Aqua Block (EastCoast Bio, North Berwick, ME) in TBS-T containing 0.1% Tween-20) for 2 hours at room temperature. Versican was detected with rabbit polyclonal antibody against mouse versican β-GAG domain (cat. no. AB1033, Millipore, Burlington, MA). Results were visualized using a LI-COR Odyssey scanner and software (LI-COR Biotechnology).

Densitometry was performed using Empiria Studio 2.3 (LI-COR Biotechnology, Lincoln, NE).

### Hyaluronan ELISA

HA quantification was assessed using a modification of reported methods. (44) Cell culture media and cell layer samples were digested with pronase (0.5 mg/ml, Roche) in 0.5 M Tris buffer (pH 6.5) for 18h at 37°C. Frozen lung tissue was minced using a blade while frozen and digested with proteinase K (0.25 mg/ml, ThermoFisher) at 60°C for 24 h. Following digestion, the pronase and proteinase K were heat-inactivated by incubation at 100°C for 20 min. HA quantity was assayed by a competitive ELISA, as previously described.(45)

### Neutrophil Adhesion Assay

Leukocyte binding was assessed using a modification of reported methods.(46) Before the adhesion assay, human promyelocytic leukemia (HL-60) cells were differentiated into neutrophil-like cells (dHL60s) by incubation of 10^6^ cells with 190µL of dimethylsulfoxide (DMSO) in 15mL of Iscove’s medium for 5 days at 37C with 5% CO_2_.(47) For adhesion on mLFs, cells were seeded in a 96-well plate and stimulated as described above. On the day of the adhesion assay, media was removed from all wells, and Hoescht stain (2µg/ml) in serum and phenol red-free RPMI medium was applied to each well and incubated for 10 minutes at 37C. mLF cell counts were quantified using live-cell fluorescent microscopy (ImageXpress Pico, Molecular Devices). After initial imaging, wells were washed once with phenol red-free RPMI.

On the day of the adhesion assay, dHL60 cells were washed three times in Gey’s medium and resuspended in the same medium at 2 x 10^6^ cells/ml. Next, dHL60s were incubated with calcein-AM (1µg/mL, Invitrogen, Carlsbad, CA) for 15 minutes at room temperature. Afterward, cells were washed twice in serum and phenol red-free RPMI, applied to mLF wells, and allowed to adhere for 60 minutes at 4°C. Cultures were washed twice with cold phenol red-free RPMI to remove non-adherent leukocytes. The adherent dHL60 cell area was quantified using live-cell fluorescent microscopy (ImageXpress Pico, Molecular Devices).

### Statistical Analysis

Statistics and images were generated using GraphPad Prism (La Jolla, CA). For PCR analyses of gene expression, normalized mRNA levels were expressed as-fold of levels in untreated or vehicle-treated controls using the comparative cycle threshold method compared with the designated housekeeping gene.(13, 48) Quantitative PCR (qPCR), BCA protein, Milliplex, and ELISA analyses were performed with two technical replicates. All data are expressed as the average ± SEM unless otherwise specified. Differences were identified by two-way ANOVA followed by Bonferroni’s posttest for multiple comparisons, with the mean of every wild-type group compared with the mean of every Col1a2/Vcan^-/-^ group with the same treatment (PBS, 3 dpi IAV, 6 dpi IAV, or 9 dpi IAV) unless otherwise specified. Statistical results with a value of *p*<0.05 were considered statistically significant.

## RESULTS

### Expression of versican and Col1a2 in C57BL/6 Mice on days 0, 3, and 6 post-IAV infection

Versican accumulates in the lungs during embryonic development, is not found in large quantities in healthy lungs, and is re-expressed in response to lung inflammation and injury during the innate immune response.(9, 13, 23, 24) The Cre-Lox P system has provided an excellent tool for the conditional deletion of versican in mice.(24, 25) For the current studies, we developed an inducible Col1a2-Cre/ERT strain of mice to delete versican in fibroblasts (i.e., Col1a2/Vcan^-/-^). To enhance the rigor of this study and better understand which cells express versican in the lungs of control mice (day 0) and in mice infected with IAV (days 3 and 6 post-infection), we utilized an existing single-cell RNA sequencing (scRNA-seq) dataset (27). This analysis provided evidence that only a select group of cells expressed versican during the 6-day course of this study (**Supplemental Fig. S1**). The highest density of versican expression observed in untreated and influenza-infected mice was in mesenchymal cells, identified by *Col1a1* and *Acta2* expression. Versican expression was also noted in epithelial cells, monocytes, neutrophils, and chondrocytes at various times during the 6-day course of the study.

To better characterize versican expression in mesenchymal cells, *Col1a1* and *Acta2* expressing clusters were isolated and reclustered. (**Fig. 1A**) Specific mesenchymal subpopulations were identified using previously described markers: alveolar fibroblasts (*Npnt*, *Ces1d*, and *Col13a1*), adventitial fibroblasts (*Pi16*, *Dcn*, and *Col14a1*), peribronchial fibroblasts (*Hhip*, *Aspn*), and smooth muscle cells (*Acta2*, *Myh11*) (**Fig. 1B**).(31, 34, 49, 50) Constitutive expression of versican was observed in all four clusters of mesenchymal cells collected from mice before infection, although variability in versican expression was noted among populations. (**Fig. 1C**). A greater proportion of adventitial fibroblasts expressed versican mRNA compared to the other two fibroblast populations (**Fig. 1C**). Adventitial and peribronchial fibroblasts showed an increase in versican expression at 3 dpi with IAV (**Fig. 1D**). Alveolar fibroblasts exhibited increased versican mRNA expression at 6 dpi with IAV. Smooth muscle cells (SMCs) demonstrated increased versican expression at 3 and 6 dpi with IAV compared to baseline (**Fig. 1D**).

**Figure 1.**
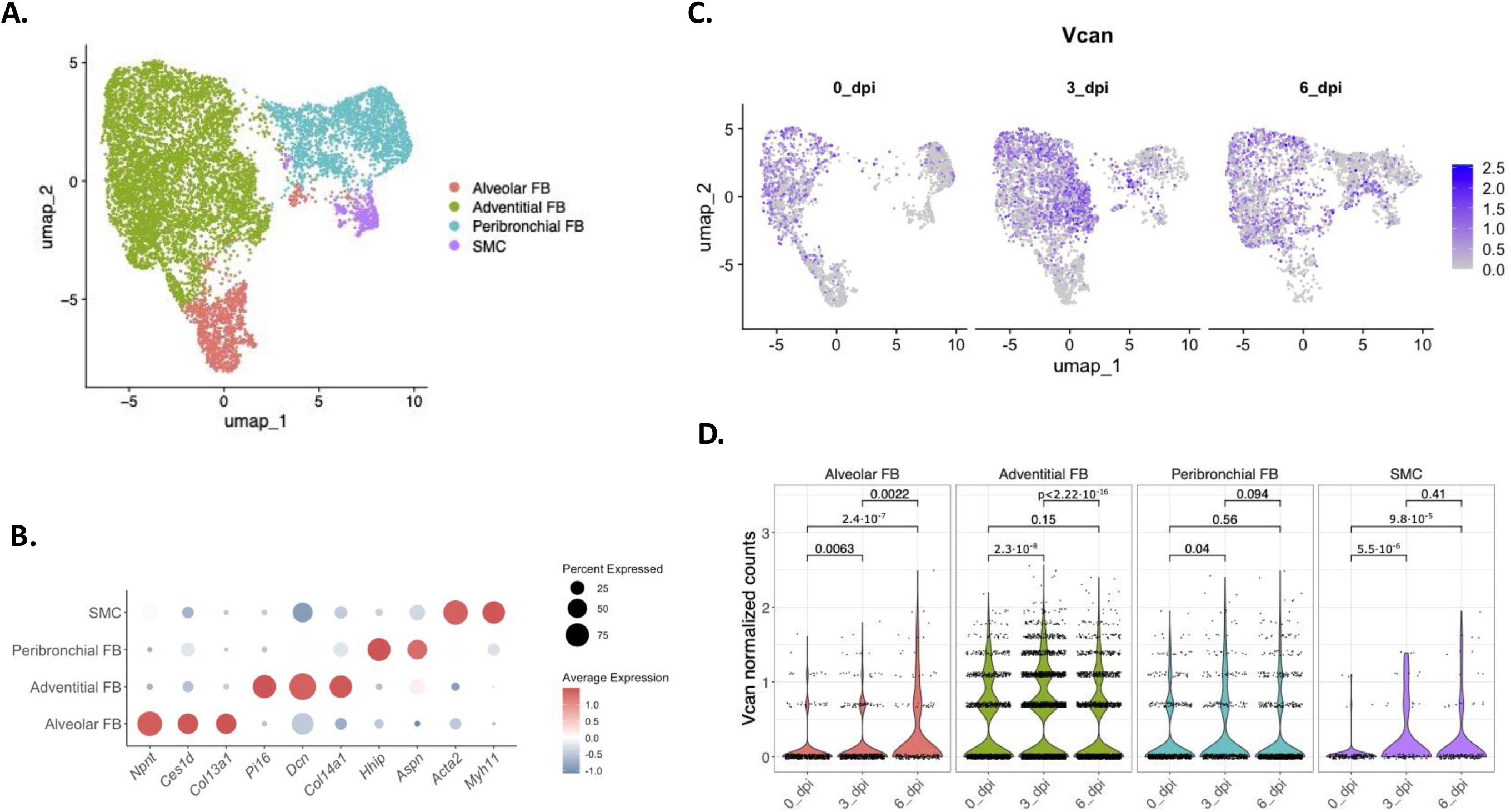
Single-cell RNA sequencing of data collected from mice 0-, 3-, and 6-dpi with IAV reveals distinct versican expression patterns in mesenchymal cells. (A) Uniform manifold approximation and projection (UMAP) visualization of mesenchymal cell clusters recovered from whole lungs of mice. (B) Dot plot showing representative markers used to define each cluster – Alveolar FBs *(Npnt, Ces1d, Col13a1)*, Adventitial FBs *(Pi16, Dcn, Col14a1)*, Peribronchial FBs *(Hhip, Aspn)*, and smooth muscle cells *(Acta2, Myh11).* (C) *Vcan* normalized counts in the four clusters of mesenchymal cells in control (0 dpi) and mice 3-and 6-dpi with IAV. (D) Statistical analysis of versican expression in alveolar FBs, adventitial FBs, peribronchial FBs, and SMCs 0-, 3-, and 6-dpi with IAV using the Wilcoxon test and p < 0.05. Abbreviations: *Vcan*, versican; SMC, smooth muscle cells; FB, fibroblasts; CPM, counts per million

The Col1a2 promoter functions as the Cre driver for tamoxifen expression, which is necessary for deleting the versican gene. To assess the specificity of Col1a2 expression, we examined its expression in the four clusters of mesenchymal cells (**Supplemental Fig. S2**). The results show that Col1a2 mRNA was present in all four clusters, with the highest density of Col1a2 expression found in alveolar and adventitial fibroblasts. This finding suggests a potential off-target effect of the Col1a2-Cre-recombinase construct, which could lead to the deletion of the versican gene in SMCs. Such an off-target effect has been recently described in cardiac SMCs.(51) The potential off-target effect complicates our ability to assert with 100% certainty that the observed effects in this study are specific to fibroblasts. As a result, we have expanded our definition of the cells undergoing genetic deletion of the versican gene to include all mesenchymal cells.

### Reduced expression and accumulation of versican in Col1a2/Vcan^-/-^ mice post-IAV infection

To better understand the extent of versican gene (*Vcan*) deletion in our versican deficient mice, *Vcan* expression was measured from whole lung homogenates of PBS-instilled (0 dpi) and IAV-instilled mice collected at 3, 6, and 9 dpi. In WT mice, peak *Vcan* expression was observed on 6 dpi (8.02 ± 1.27, fold change relative to WT uninjured), consistent with previously published findings in C57BL/6J mice (**Fig. 2A**).(23) Col1a2/Vcan^-/-^ mice had significantly less *Vcan* expression on 6 dpi (3.91 ± 0.56, fold change relative to WT uninjured). There were no statistically significant differences in the amount of *Vcan* expressed between Col1a2/Vcan^-/-^ and WT at 3-and 9-dpi, however 9-dpi demonstrated a trend for decreased *Vcan* expression (p = 0.051). These data demonstrated the successful knockdown of *Vcan* in our mouse model and that mesenchymal cells, particularly fibroblasts, are a significant source of *Vcan* expression in the lungs during IAV infection.

**Figure 2.**
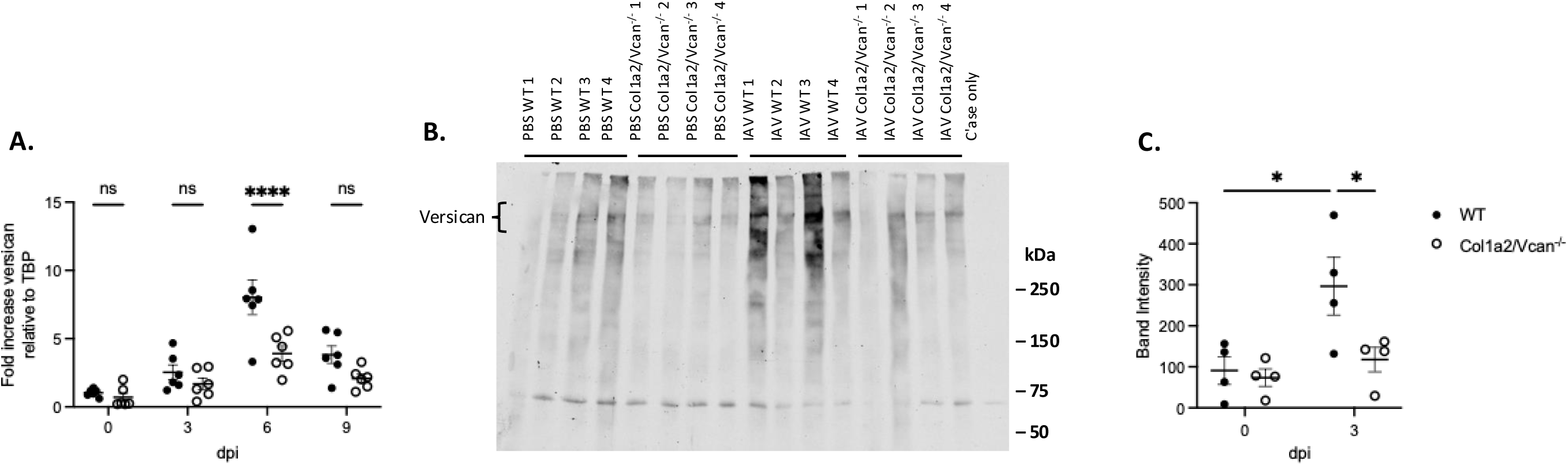
Expression and synthesis of versican in the lungs post-IAV infection in mice deficient in fibroblast-derived versican. (A) mRNA was collected from lung homogenates at 0 (PBS), 3, 6, and 9 dpi with IAV. Quantitative real-time PCR was used to determine the fold change in versican gene expression relative to WT uninjured. mRNA expression was normalized to TBP. (B) Western blot of lung tissue from WT and Col1a2/Vcan^-/-^ mice treated with PBS or IAV at 3 dpi, were incubated with a rabbit anti-mouse antibody targeting the beta-glycosaminoglycan binding domain within versican. (C) The data in (B) were quantified using densitometry with ImageJ software. Values are mean ± SEM, *n* = 6 in (A) and *n* = 4 in (C). Two-way ANOVA with Bonferroni’s for multiple comparison test \**p*≤0.05, \*\*\*\**p*≤0.0001 for (A) and (C). Abbreviations: WT, wild-type; Col1a2/Vcan^-/-^, mice deficient in mesenchymal-derived versican; IAV, influenza A virus; dpi, days post-infection; TBP, TATA-box binding protein

To assess the knockdown of versican protein in Col1a2/Vcan^-/-^ mice during the innate immune response to IAV infection, versican accumulation in IAV-instilled lungs on 3 dpi was measured by Western blot (**Fig. 2B**). An antibody targeting the beta-glycosaminoglycan (β-GAG) binding domain was used to determine the presence of V0 and V1 versican isoforms.(20) V0 and V1 are the predominant versican isoforms in the lungs during homeostasis and under inflammatory conditions.(9, 24) In both Col1a2/Vcan^-/-^ and WT IAV-infected mice, bands corresponding to V0 and V1 at ∼550 kDa and ∼500 kDa, respectively, were present (**Fig. 2B**). Additionally, smaller fragments with distinct bands were observed, consistent with previous publications.(24, 43) Densitometry analysis showed a significant increase in band intensities for V0/V1 versican in IAV-infected WT mice compared to PBS-instilled WT mice (**Fig. 2C**). The mean band intensity was compared, and IAV-infected WT mice had a 225% increase in the amount of versican compared to PBS-instilled WT mice. Additionally, there was a significant decrease in versican band intensity for IAV-infected Col1a2/Vcan^-/-^ mice compared to IAV-infected WT mice at 3 dpi (**Fig. 2C)**. Col1a2/Vcan^-/-^ mice had a 60% decrease in versican accumulation compared to WT mice. These data indicated successful versican protein depletion in our mouse model and that mesenchymal cells are a significant source of versican in the lungs during IAV infection.

### Influenza disease severity and viral load were unaltered throughout IAV infection in Col1a2/Vcan^-/-^ mice

Col1a2/Vcan^-/-^ and WT mice were infected with 20 PFU of IAV and monitored for up to 15 dpi to investigate the impact of mesenchymal-derived versican deficiency on influenza infection and disease. This dose of IAV caused severe influenza pneumonia in which mice approached humane euthanasia endpoint criteria but ultimately survived infection.(23) Changes in body weight and lung viral copy numbers were measured. Both Col1a2/Vcan^-/-^ and WT mice exhibited similar body weight loss and recovery profiles; the peak loss of ∼25% of initial body weight reflected the marked severity of disease with this high dose of IAV in both strains of mice.(23, 38) (**Fig. 3A**). In both Col1a2/Vcan^-/-^ and WT mice, viral copies in whole lung homogenates peaked at 6 dpi, with no significant differences observed between mouse strains at any time point before viral clearance (**Fig. 3B**). These data suggested that mesenchymal-derived versican is a component of the host immune response to IAV infection that does not directly impact IAV proliferation or clearance in the lungs.

**Figure 3.**
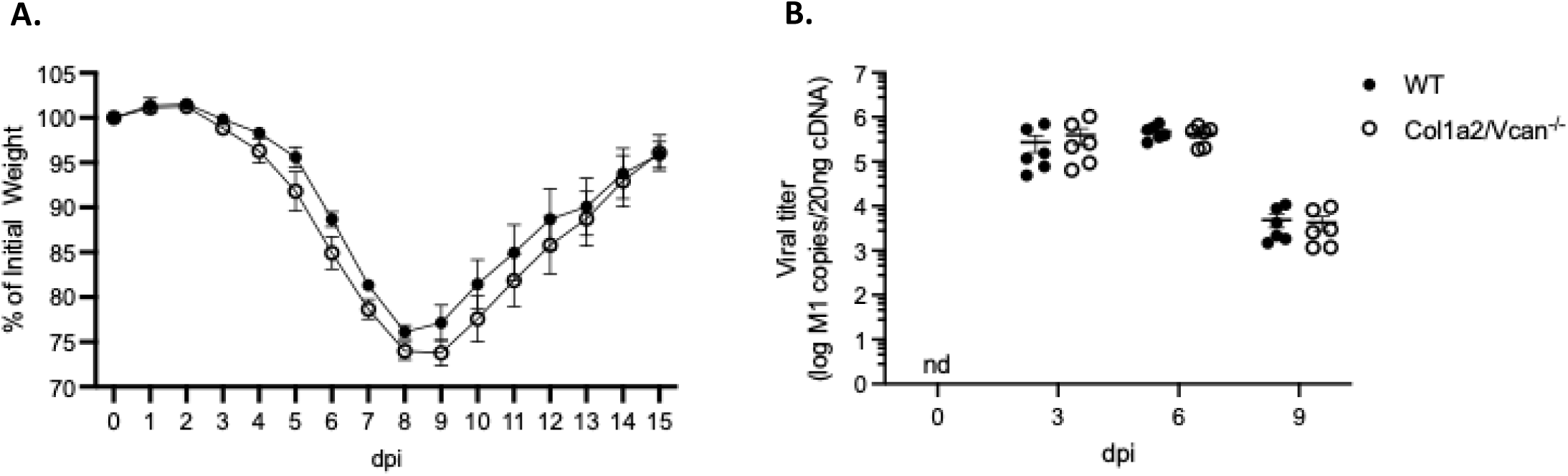
Disease severity and viral load during IAV infection. (A) Mouse percentage of initial body weight following infection with IAV over 15 dpi. Both WT and Col1a2/Vcan^-/-^ mice reached their maximum weight loss by 9 dpi. (B) mRNA was collected from lung homogenates at 0 (PBS), 3, 6, and 9 dpi with IAV and quantitative real-time PCR was used to determine the absolute number of viral copies of the *M1* gene of IAV. DNA from purified plasmid was used to generate a standard curve for copy number determination. Values are mean ± SEM, *n* = 4 in (A) and *n* =6 in (B). A repeated measures two-way ANOVA with Bonferroni’s for multiple comparison test for (A) and two-way ANOVA with Bonferroni’s for multiple comparison test for (B). Abbreviations: nd, not detected

### Reduced inflammatory cell recruitment in Col1a2/Vcan^-/-^ mice infected with IAV

To test our hypothesis that mesenchymal-derived versican is pro-inflammatory and enhances the innate immune response to IAV infection, we assessed several metrics of acute lung injury in Col1a2/Vcan^-/-^ and WT mice. Acute lung injury (ALI) is a multidimensional entity characterized by key pathophysiologic features that the American Thoracic Society has categorized into four domains, which include 1) histological evidence of tissue injury, 2) alterations of the alveolar-capillary barrier, 3) presence of an inflammatory response, and 4) physiologic dysfunction.(52) First, our studies looked for differences in histologic injury and permeability of the alveolar-capillary barrier between Col1a2/Vcan^-/-^ and WT mice. In response to oropharyngeal exposure to 20 PFU of IAV, we observed severe injury as indicated by the percent of lungs involved and the severe degree of inflammation, edema, and hemorrhage. Semiquantitative histological analysis of lung inflammation and injury showed no differences in pulmonary injury between Col1a2/Vcan^-/-^ and WT mice at 9 dpi with 20 PFU of IAV (**Supplemental Fig. S3A**).

Assessment of the alveolar-capillary barrier by measurement of total protein present in the BAL throughout infection revealed no significant differences between Col1a2/Vcan^-/-^ and WT mice up to 9 dpi with IAV (**Supplemental Fig. S3B**). However, BAL total protein was markedly elevated from baseline after IAV infection, reflecting the severity of injury and loss of alveolar-capillary barrier integrity on both strains.

Next, spectral flow cytometry was used to assess differences in inflammatory responses between Col1a2/Vcan^-/-^ and WT mice up to 6 dpi with IAV. The spectral flow cytometry panel applied was broad, immunophenotyping > 90% of all leukocytes present (**Table 1**).(39) This allowed for an unbiased and exploratory investigation of inflammatory leukocyte recruitment into the lungs.(39) Additionally, *in vivo* labeling of intravascular leukocytes was included in the spectral flow cytometry panel, which allowed for the assessment of leukocyte recovery from 3 distinct pulmonary microenvironments – airways, lungs, and pulmonary vasculature. Cells measured in the airways were those collected by bronchoalveolar lavage. Cells measured in the lungs were collected from single-cell suspensions of lung homogenates that were negative for the intravascular leukocyte label (CD45IV^-^). Vasculature cells were CD45IV^+^ cells collected from the single-cell suspensions of lung tissue. Vasculature cells remained within the pulmonary vessels after perfusion of the systemic vasculature and were potentially poised to enter the lung interstitium.(39)

During early IAV infection, neutrophil, monocyte, and dendritic cell recruitment into the airways was significantly reduced in Col1a2/Vcan^-/-^ mice at 3 dpi with IAV. Col1a2/Vcan^-/-^ mice had 80% fewer neutrophils, 64% fewer monocytes, and 67% fewer dendritic cells in the airways on 3 dpi than WT mice (**Fig. 4A**). The recruitment of neutrophils and monocytes into the lungs was also significantly attenuated at 3 dpi with IAV, while dendritic cell recruitment to this compartment had a trend toward significant attenuation in Col1a2/Vcan^-/-^ mice compared to WT mice. In the lungs, Col1a2/Vcan^-/-^ mice had 75% fewer neutrophils and 47% fewer monocytes compared to WT mice (**Fig. 4B**). Finally, only neutrophils were reduced in number in the vasculature of Col1a2/Vcan^-/-^ mice on 3 dpi, with 45% fewer neutrophils than wild-type mice (**Fig. 4C**). No differences were found for other leukocyte populations. These data suggest that mesenchymal-derived versican plays a role in neutrophil, monocyte, and dendritic cell recruitment to the lungs during the innate immune response to IAV infection.

**Figure 4.**
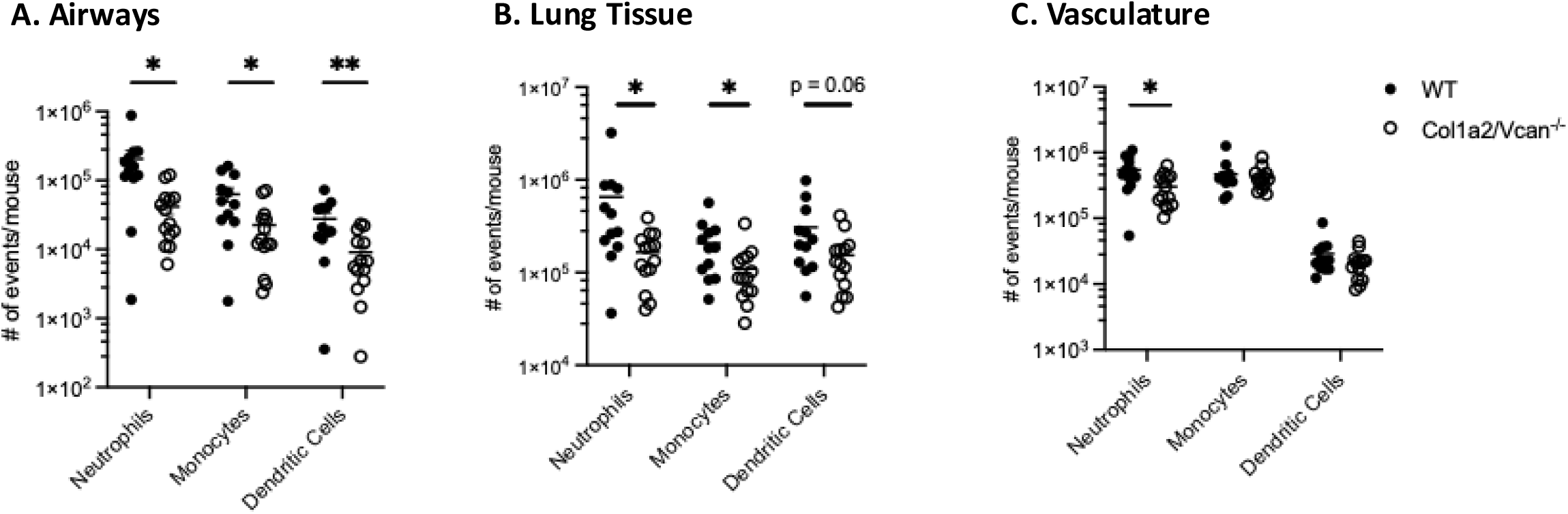
Neutrophil, monocyte, and dendritic cell recruitment to pulmonary microenvironments on 3 dpi with IAV. Neutrophil, monocyte, and dendritic cell recruitment to the airways (A), lungs (B), and marginated vasculature (C) was determined in Col1a2/Vcan^-/-^ and WT mice 3 dpi with IAV. Col1a2/Vcan^-/-^ mice demonstrated reduced numbers of neutrophils across all three microenvironments compared to WT mice. Monocytes were reduced in the airways and lungs of Col1a2/Vcan^-/-^ mice, while dendritic cells were only significantly reduced in the airways. Values are mean ± SEM, *n* = 12-14 in (A), (B), and (C). Student’s unpaired t-test \**p*≤0.05, \*\**p*≤0.01 for (A), (B), and (C).

Neutrophil, monocyte, and dendritic cell populations mobilize from the bone marrow during infection. Therefore, it was necessary to determine whether Col1a2/Vcan^-/-^ mice had adequate myelopoiesis during IAV infection. To investigate the possibility of a defect in myelopoiesis, the spectral flow cytometry panel was used to analyze whole blood samples from mice 3 dpi with IAV. No differences in circulating leukocyte populations were found between Col1a2/Vcan^-/-^ and WT mice (**Supplemental Fig. S4)**. Leukocyte recruitment to the lungs was also investigated at 6 dpi with IAV. At this time, Col1a2/Vcan^-/-^ mice had fewer mono-macrophages than WT mice (**Fig. 5**). Col1a2/Vcan^-/-^ mice had 45% and 37% fewer mono-macrophages in the vasculature and lungs, respectively (**Fig. 5**). All other leukocyte populations were comparable between Col1a2/Vcan^-/-^ and WT mice in each microenvironment investigated at 6 dpi (not shown). These data support the role of mesenchymal-derived versican in neutrophil migration from the circulation to margination within the pulmonary vasculature and, subsequently, neutrophil migration into the lungs and airways. Additionally, these data support the role of mesenchymal-derived versican in monocyte and dendritic cell migration into the airways. Taken together, the consequences of reduced neutrophil and monocyte migration in Col1a2/ Vcan^-/-^ mice were not reflected in other ALI measures but suggested that other aspects of the inflammatory response may be altered in Col1a2/Vcan^-/-^ mice compared to WT mice.

**Figure 5.**
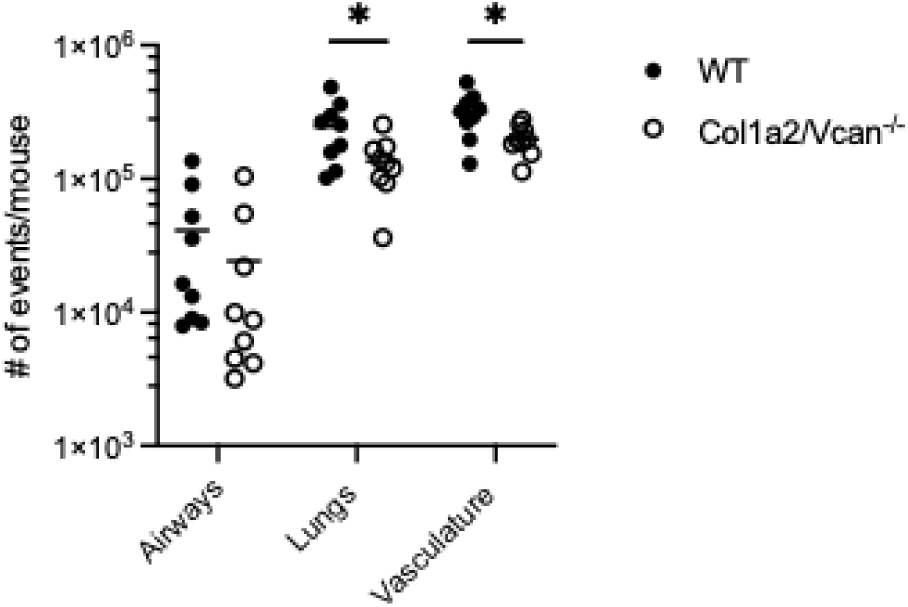
Mono-macrophage recruitment to pulmonary microenvironments on 6 dpi with IAV. Mono-macrophage recruitment to the vasculature, lungs, and airways was determined in Col1a2/Vcan^-/-^ and WT mice 6 dpi with IAV. Col1a2/Vcan^-/-^ mice demonstrated reduced numbers of mono-macrophages in the marginated vasculature and lungs compared to WT mice. Values are mean ± SEM, *n* = 9, Student’s unpaired t-test \**p*≤0.05

### Unaltered production of inflammatory cytokines and chemokines in Col1a2/Vcan^-/-^ mice post-IAV infection

Previous studies have reported alterations in chemokine and cytokine production in versican deficient mice exposed to PolyI:C and RSV.(24–26) Therefore, the presence of soluble inflammatory mediators was evaluated to understand how the inflammatory response in Col1a2/Vcan^-/-^ mice may be altered during IAV-induced acute lung injury. Concentrations of inflammatory mediators were determined in the BAL fluid from PBS-instilled (0 dpi) and IAV-instilled Col1a2/Vcan^-/-^ and WT mice at 3 and 6 dpi using a mouse immunology multiplex magnetic bead assay. The multiplex assay measured IL-4, IL-6, IL-10, IL-13, CCL2, CXCL1, CXCL2, TNFα, and VEGF. No significant differences between mouse strains were detected at any time for any inflammatory mediator (**Fig. 6A-I**). Neutrophils had the greatest reduction across all microenvironments evaluated of the leukocytes with reduced recruitment in Col1a2/Vcan^-/-^ mice (**Fig. 4**). Therefore, additional effort was made to evaluate neutrophil chemokine gradients for CXCL1 and CXCL2. Plasma, lung homogenates, and BAL fluid samples were collected from Col1a2/Vcan^-/-^ and WT mice 3 dpi and assessed by ELISA. No significant differences in CXCL1 and CXCL2 concentrations were observed between Col1a2/Vcan^-/-^ and WT mice (**Fig. 7A-B**). These data suggested that the reduced recruitment of leukocytes into the lungs of Col1a2/Vcan^-/-^ mice was not due to chemokine or cytokine-mediated mechanisms.

**Figure 6.**
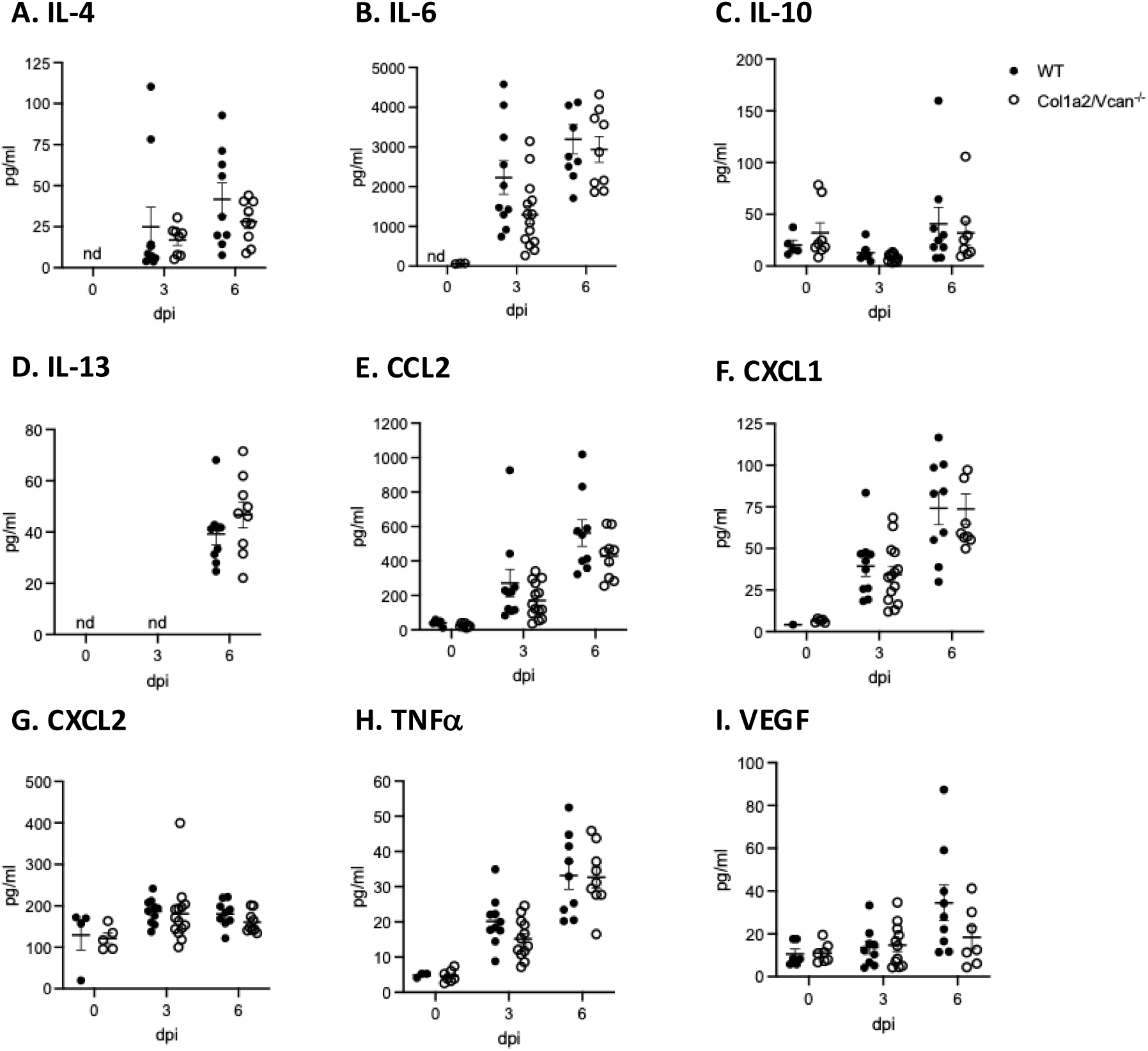
Concentrations of chemokines and cytokines from BAL fluid during IAV. (A-I) Values are mean ± SEM with *n*= 6-14 per time point. Groups were statistically tested using two-way ANOVA with Bonferroni’s for multiple comparison test.

**Figure 7.**
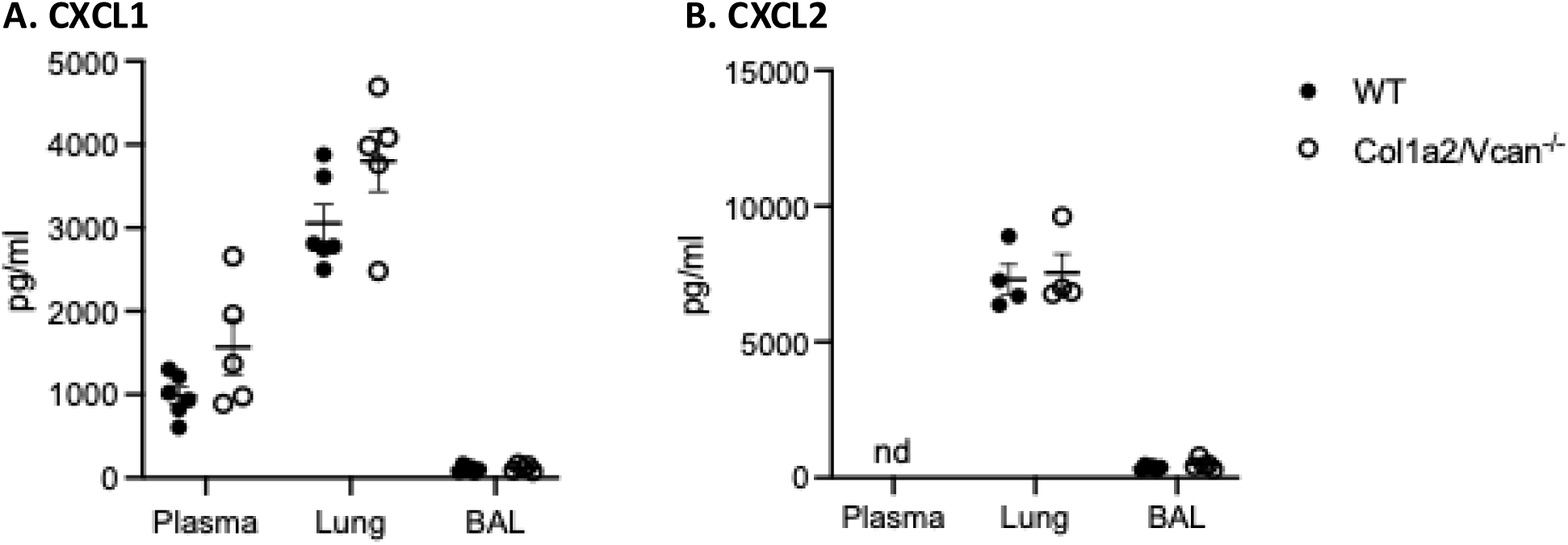
Concentrations of CXCL1 and CXCL2 from paired plasma, lung, and BAL fluid 3 dpi with IAV. (A) CXC1 and (B) CXCL2 were measured by ELISA in plasma, lung, and BAL samples collected from Col1a2/Vcan^-/-^ and WT mice 3 dpi with IAV. Values are mean ± SEM with *n*=4-6 per time point. Groups were statistically tested using two-way ANOVA with Bonferroni’s for multiple comparison test. Abbreviations: BAL, bronchoalveolar lavage; dpi, days post infection; IAV, influenza A virus.

### Reduced binding of neutrophils to the ECM of Col1a2/Vcan^-/-^ mouse lung fibroblasts

Since the profile of CXCL1 and CXCL2 in BAL fluid and plasma appeared similar between Col1a2/Vcan^-/-^ and WT mice, we investigated whether the reduction in leukocyte recruitment to the lungs of IAV-infected Col1a2/Vcan^-/-^ mice was due to reduced leukocyte adhesion to the ECM of lung fibroblasts, using a well-established *in vitro* adhesion assay.(21, 24) We specifically investigated neutrophil adhesion as previous studies have demonstrated that monocytes bind to versican-enriched matrices produced by fibroblasts and have reduced adhesion to versican-deficient fibroblasts.(17, 24, 53) Fibroblasts cultured from the lungs of Col1a2/Vcan^-/-^ and WT mice were stimulated with poly(I:C) to generate a versican and hyaluronan-enriched matrix, as previously described.(17, 24, 53) A representative western blot for versican isoform V0/V1 collected from cultured mouse lung fibroblasts (mLFs) is shown in **Fig. 8A**. Col1a2/Vcan^-/-^ mLFs had a visible knockdown of V0/V1 (∼550-500 kDa) when treated with poly(I:C) compared to WT mLFs. To determine whether versican was important for neutrophil adhesion to lung fibroblasts, we assessed the adhesion of differentiated human promyelocytic leukemia (dHL60) cells to Col1a2/Vcan^-/-^ and WT mLFs stimulated with poly(I:C).(21, 47) This *in vitro* adhesion model was adapted from published studies that utilized mouse-derived stromal cells with human leukocyte cell lines and was chosen for our experiments as it allowed us to utilize our novel Col1a2/Vcan^-/-^ mLFs.(21, 24, 54) We found that dHL60s were significantly less adherent to Col1a2/Vcan^-/-^ mLFs compared to WT mLFs, whether unstimulated or stimulated with poly(I:C) (**Fig. 8B**). These findings suggest that versican may be critical for the adhesion of neutrophils to fibroblasts and other mesenchymal cells as they move from the vasculature to the airways through the pulmonary interstitial space.

**Figure 8.**
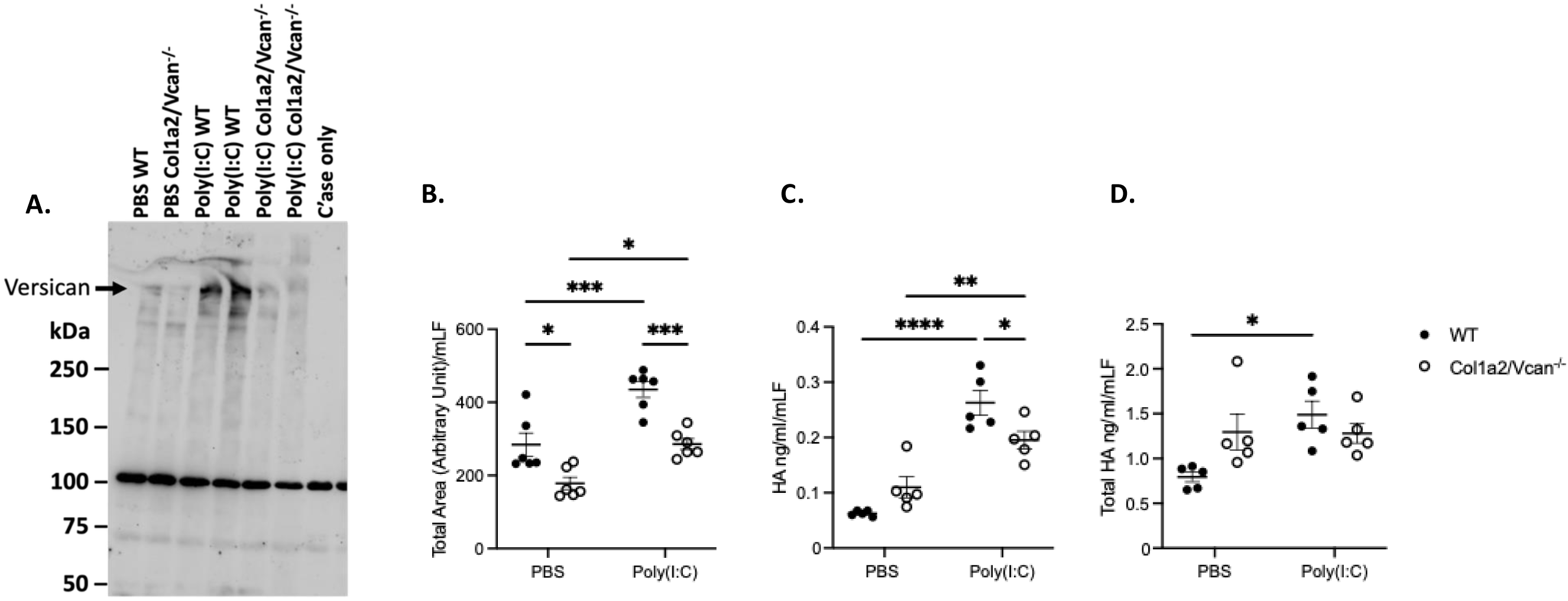
Versican protein, cell-associated hyaluronan, and neutrophil adhesion in mLFs stimulated with Poly(I:C) (A) Protein of versican V0/V1 (∼550-500 kDa, black arrow) in mLFs from Col1a2/Vcan^-/-^ and WT mice stimulated with poly(I:C) measured by Western blot. (B) Adhesion of dHL60s to mLFs from Col1a2/Vcan^-/-^ and WT mice stimulated with poly(I:C) were quantified using live-cell fluorescent microscopy and normalized to mLFs cell counts. (C) Cell-associated hyaluronan measured by ELISA in mLFs stimulated with poly(I:C) from Col1a2/Vcan^-/-^ and WT mice. (D) Cell-associated and cell culture media hyaluronan measured by ELISA in mLFs stimulated with poly(I:C) from Col1a2/Vcan^-/-^ and WT mice. Values are mean ± SEM with *n*=5. Asterisks show groups that are statistically significantly different (\**p*≤0.05, \*\**p*≤0.01, \*\*\**p*≤0.001, \*\*\*\**p*≤0.0001) using two-way ANOVA with Bonferroni’s for multiple comparison test. Abbreviations: mLFs, mouse lung fibroblasts; Poly(I:C), polyinosinic:polycytidylic acid; dHL60s, differentiated HL60 cells; C’ase only, chondroitinase ABC enzyme buffer control

Hyaluronan (HA) is a binding partner of versican and plays an important role in leukocyte adhesion in the lungs of IAV-infected mice.(21) In previous studies, treatment of WT lung fibroblasts with poly(I:C) *in vitro* resulted in increased cell-associated HA, a corresponding reduction of HA in the culture medium, and the colocalization of HA with versican.(17, 53) When mLFs from mice with a global deficiency in versican were stimulated with poly(I:C), cell-associated HA cable structures were not observed, suggesting that versican deficiency disrupts HA cable formation.(24) To investigate the content of HA in our culture system, HA ELISA was performed on mLF cell layers after Col1a2/Vcan^-/-^ and WT mLFs were stimulated with poly(I:C). We found WT mLFs had significantly increased cell-associated HA when stimulated with poly(I:C) compared to PBS-treated WT mLFs (**Fig. 8C**). Additionally, when stimulated with poly(I:C) Col1a2Vcan^-/-^ mLFs retained less cell-associated HA than WT cells (**Fig. 8C**). To investigate whether the change in HA accumulation on mLFs was due to changes in HA synthesis, the amount of HA in the culture media was quantified by ELISA. There was no significant difference in the total amount of HA (cell-associated + medium) between WT and Col1a2Vcan^-/-^ mLFs stimulated with poly(I:C) (**Fig. 8D**). Interestingly, Col1a2Vcan^-/-^ mLFs had a trend for higher total levels of HA (cell-associated + medium) than WT mLFs when mLFs were treated with PBS and did not increase the total amount of HA in response to poly(I:C) like WT mLFs did. However, there was significantly increased cell-associated HA in Col1a2Vcan^-/-^ mLFs stimulated with poly(I:C) compared to PBS-treated Col1a2Vcan^-/-^ mLFs (**Fig. 8C**). In summary, our results suggested that the observed reduction of cell-associated HA in versican-deficient cultures was due to the absence of versican-mediated retention of HA on the cell surface of lung fibroblasts. Taken together, our data support that versican, in conjunction with cell-associated hyaluronan, is important for neutrophil adhesion to mouse lung fibroblasts and that adhesion to pulmonary fibroblasts may be a mechanism by which versican facilitates neutrophil emigration into the lungs during IAV infection *in vivo*.

## DISCUSSION

Consistent with the study objectives, we determined the time course of changes in mesenchymal-derived versican expression and the impact of mesenchymal-derived versican deficiency on leukocyte migration into pulmonary and vascular microenvironments during IAV infection, which have not been previously reported. In addition, we document the following significant findings: (1) adventitial fibroblasts have moderate constitutive expression of versican in uninfected mice and increase their expression of versican on 3 dpi (2) neutrophils in the airways, lungs, and marginated pulmonary vasculature are significantly reduced in Col1a2/Vcan^-/-^ mice compared to WT controls on 3 dpi with IAV; (3) monocytes in the airways and lungs are significantly reduced on 3 dpi with IAV, and this is followed by significantly reduced numbers of mono-macrophages on 6 dpi in Col1a2/Vcan^-/-^ mice compared to WT controls; and (4) a differentiated neutrophil-cell line is less adherent to versican-deficient fibroblasts and versican-deficient fibroblasts have significantly reduced cell-associated hyaluronan (HA) content *in vitro*. These findings suggest that mesenchymal-derived versican and cell-associated HA are necessary for the adhesion of neutrophils and monocytes to lung fibroblasts. This adhesion likely occurs as they transit from the pulmonary vasculature across the endothelial barrier through the pulmonary interstitium. Neutrophils subsequently transit from the interstitium across the respiratory epithelium into the airways of mice infected with influenza A virus. Additionally, we did not find evidence of neutrophil or monocyte recruitment attenuation by mesenchymal-derived versican through chemokine or cytokine-mediated mechanisms. These findings emphasize the importance of mesenchymal-derived versican in generating specialized provisional matrices within distinct microenvironments as part of the host immune response to IAV infection.

The moderate constitutive expression of versican by fibroblasts in uninfected mice (0 dpi) was not anticipated based on our previous studies showing minimal accumulation of versican protein in healthy lungs.(9, 23, 24) However, the work of Potter-Perigo et al. demonstrates that human fibroblasts constitutively express versican *in vitro*, and that the increase in versican synthesis in human fibroblasts treated with poly(I:C) was not due to heightened expression of versican, but rather through post-transcriptional regulation and reduction in the proteases that degrade versican, such as ADAMTS-1,-4, and-5.(17) Our study also supports the post-transcriptional control of versican in healthy lungs as the amount of versican accumulation from PBS-instilled lungs was much less than that seen at 3 dpi by Western blot. Future studies will need to address the mechanisms regulating the expression and accumulation of versican by fibroblasts in the lungs of healthy mice.

During IAV infection, mesenchymal-derived versican plays an important contextual role in regulating leukocyte migration in distinct pulmonary and vascular microenvironments.

Interestingly, neutrophil migration into the lungs and airways is reduced by 75% and 80% in Col1a2/Vcan-/-mice during peak neutrophil emigration from the vasculature. Additionally, neutrophils are reduced in the marginated vasculature of Col1a2/Vcan-/-mice on 3 dpi by 45%, but there is no difference in circulating neutrophil numbers between Col1a2/Vcan-/-mice and WT controls. Thus, we interpret that neutrophils are less adherent to the pulmonary endothelium in the Col1a2/Vcan-/-mice. Additionally, our finding suggests neutrophils are less adherent to versican-deficient fibroblasts. Several adhesion molecules, including CD44, L-selectin, PSGL-1, and β-integrin, are known to bind versican.(21) Future studies will need to explore whether mesenchymal-derived versican influences endothelial phenotype or function, and whether interactions between specific adhesion molecules and versican are necessary for neutrophil adhesion and migration into the lungs of mice infected with IAV.

Naive and inflamed lungs possess a neutrophil reservoir with 40-65 times more neutrophils sequestered in the pulmonary capillaries than in larger vessels.(55) This is primarily due to the mechanical trapping of neutrophils in pulmonary capillaries, which occurs independently of adhesion mechanisms initiated by selectins and integrins.(55, 56) Under inflammatory conditions, the role of selectins and integrins in neutrophil adhesion to the pulmonary endothelium and recruitment into the lungs has been shown to vary with the type of inflammatory stimuli.(56–58) Neutrophil migration was dependent on CD11/CD18 in mice infected with gram-negative pneumonia, while CD11/CD18 independent emigration was observed in mice infected with *S. pneumoniae*.(57, 58) While it is unknown whether neutrophil adhesion and emigration during IAV infection occur via integrin or selectin dependent or independent processes, in our study, neutrophils in the pulmonary marginated vasculature are those cells resistant to vascular perfusion. Soucy et al. demonstrated that in mice infected with *S. pneumoniae*, alveolar fibroblast-derived versican was not necessary for neutrophil migration into the lungs.(30) Together with our findings, this raises important questions about whether versican-dependent or independent emigration of leukocytes into the lungs of mice varies according to the type of inflammatory stimuli and/or the specific fibroblast subtype producing versican.

To investigate neutrophil adhesion to versican-deficient fibroblasts, we performed adhesion assays with mouse lung fibroblasts (mLFs) from Col1a2/Vcan^-/-^ mice and WT controls. We found that versican-deficient mLFs had significantly reduced adhesion of dHL60 neutrophil-like cells after stimulation with poly(I:C), a TLR-3 agonist that generates a versican-enriched ECM in WT cells.(53) Previous studies have shown that monocytes have reduced adhesion to versican-deficient fibroblasts, however, these are novel findings for a neutrophil cell line.(24) Additionally, we demonstrated that versican-deficient fibroblasts have reduced cell-associated HA compared to WT fibroblasts after poly(I:C) stimulation. To our knowledge, these results are the first reported quantification of cell-associated HA in versican-deficient mLFs. Our findings align with previous studies and suggest that versican plays an important role in sequestering cell-associated HA within provisional matrices. (17, 24, 53) Future studies will need to address the adhesion molecules on neutrophils and monocytes required for their adhesion to the cell-associated versican-HA matrices formed by fibroblasts.

Whether versican, HA, or both are required for neutrophil adhesion remains unknown. Experiments applying versican antibody or hyaluronidase to WT mLFs have the potential to clarify this point.(59) Tang et al. demonstrated that dHL60 cells did not have increased adherence *in vitro* to immobilized HA alone but depended on the presence of heavy chains (HCs) and the formation of an HC•HA matrix produced by WT mLFs. The HC•HA matrix is produced by a TNFα-stimulated gene-6 (TSG-6) dependent process by transferring HCs from the inter-alpha-trypsin inhibitor (IαI) to HA. It is difficult to interpret how versican may have contributed to dHL60 adhesion in these studies. Additionally, it is unknown what role IαI and HCs may have played in dHL60 adhesion to matrices deficient in versican in our studies. We utilized DMEM with 2% FBS as the medium to treat our cells with poly(I:C). Serum is a source of IαI and would have donated HCs to the HA based on our previous work.(21) *In vivo,* it has been previously shown that HA accumulates in the BAL fluid and lungs post-IAV infection.(21, 60) However, we did not observe a significant reduction of HA in BAL fluid or whole lung homogenates from IAV-infected Col1a2Vcan^-/-^ mice compared to WT mice (not shown). This could be due to other cellular sources of HA, such as HA derived from epithelial cells, endothelial cells, pericytes, and smooth muscle cells, obscuring potential differences in the HA content associated with the ECM of fibroblasts in our model. Future work is needed to determine the specific contribution of HA, IαI, and HCs to the adhesion of neutrophils on versican-enriched matrices *in vitro* and *in vivo*.

Similar to neutrophils, we found that mesenchymal-derived versican plays a vital role in regulating monocyte migration in the lungs and airspaces during IAV infection. Monocyte migration into the lungs and airways is reduced by 47% and 64%, respectively, in Col1a2/Vcan^-/-^ mice compared to WT controls. We did not observe differences in monocytes in the marginated vasculature as we did with neutrophils and, therefore, interpret that monocytes are adhering to the pulmonary endothelium during IAV infection equivalently between each mouse strain. This may suggest that monocytes interact with different adhesion molecules than neutrophils and, therefore, are less dependent on the mesenchymal-derived versican-HA matrix to emigrate into the lungs from the vasculature. However, adhesion to lung fibroblasts is likely important for both cell types for emigration into the airways. The monocyte cell line, U937, has previously been shown to have reduced adhesion to versican-deficient mLFs stimulated with poly(I:C).(24) The reduction of monocytes into the lungs and airways on 3 dpi with IAV may also contribute to the observed reductions in monocyte-derived cell populations, such as the 67% reduction of airway dendritic cells on 3 dpi and 45% reduction of lung mono-macrophages observed shortly after on 6 dpi. Additionally, monocyte and neutrophil trafficking into the lung are interdependent, with monocytes facilitating neutrophil transendothelial migration from the vasculature into the lungs.(56, 61) Thus, in addition to the decreased adhesion of neutrophils to versican-deficient fibroblast, the reduced number of monocytes in the lungs also likely contributes to the reduction in neutrophils observed on 3 dpi in Col1a2/Vcan^-/-^ mice.

Macrophages are resident phagocytic cells localized in tissues, and our spectral flow cytometry panel was compatible with immunophenotyping for alveolar macrophages, recruited macrophages, and mono-macrophages. However, macrophages were not expected to localize to vascular microenvironments (IVCD45^+^ macrophages), and identifying mono-macrophages in the marginated vasculature was unexpected (**Fig. 5**). Several possible explanations exist for mono-macrophage localization in the marginated vasculature. One explanation is that IVCD45^+^ labeled mono-macrophages were monocytes that had become adherent to the pulmonary vasculature and increased their expression of F4/80 and CD64 while migrating from the vasculature into the pulmonary interstitium. The decreased adhesion of monocytes to the pulmonary endothelium that transition to mono-macrophages as they migrate into the lungs could explain the reduced number of mono-macrophages recovered from the marginated vasculature and lungs in Col1a2/Vcan^-/-^ mice. A second possibility is that these IVCD45^+^ labeled mono-macrophages are monocytes that became adherent to pulmonary endothelial cells and remain in the pulmonary circulation as macrophages. Work from several groups shows that rats exposed to 0111:B4 lipopolysaccharide (LPS) intravenously or intraperitoneally have a significant but transient increase in activated phagocytic mononuclear cells in the pulmonary microvasculature.(62, 63) This monocyte population has been referred to as induced pulmonary intravascular macrophages since they take on some but not all the characteristics of the pulmonary intravascular macrophages found in sheep, goats, pigs, and other species.(64, 65) Finally, we cannot rule out the possibility that the IVCD45^+^ cells are interstitial cells that were identified as being in the vasculature due to increased vascular permeability. Future studies will need to determine the origin of the IVCD45^+^ macrophages identified in this study.

A limitation of our studies can be attributed to the employment of our spectral flow cytometry panel to assess circulating leukocytes as it was optimized for identifying CD11b^+^ and CD103^+^ conventional dendritic cells (cDCs) in the airways and lung tissue and, therefore, it lacked the markers necessary to identify precursor-cDCs. Precursor-cDCs migrate from the bone marrow through the vasculature to the lungs in response to CCR2 ligands and differentiate into cDCs within the lungs during inflammation.(66, 67) Therefore, we did not determine if there was a difference in circulating precursor-cDCs between Col1a2/Vcan^-/-^ and WT mice. However, no difference was found in total circulating leukocytes or other myeloid-lineage cells (**Supplemental Fig. S4**). Some inflammatory dendritic cells are monocyte-derived; therefore, the observed reduction in dendritic cells may be partly due to the concurrent decrease in monocytes in the lungs and airways on 3 dpi.(68)

Our studies used several methodologies to look for evidence of differences in disease and lung injury between Col1a2/Vcan^-/-^ and WT mice. Mice from both strains had similar weight loss during IAV infection. Additionally, no significant differences were seen in total protein levels in the BAL fluid throughout infection. Acute lung injury scores did not reveal differences between mouse strains post-IAV infection. The contribution of neutrophils to lung injury during IAV infection is complex. Existing literature supports both a protective role for neutrophils and a role for excessive neutrophil activity in contributing to severe clinical disease.(69) A potential limitation of our studies is that we chose a high dose of IAV, 40% of the euthanasia dose 50, which resulted in severe lung injury and clinical disease. Differences in acute lung injury and clinical disease between Col1a2Vcan^-/-^ and WT mice may be elucidated in studies utilizing a lower dose of IAV or models of neutrophil-dependent lung injury from such as *Pseudomonas aeruginosa*.(23, 38, 70) Another potential limitation of our study’s assessment of IAV-induced lung injury is that we only investigated histologic injury up to 9 dpi in our model. The differences in early neutrophil and monocyte recruitment to the lungs could expedite the resolution of IAV-induced lung injury further in the time course of the disease process in Col1a2/Vcan^-/-^ mice. Future studies are needed to investigate this possibility.

In conclusion, our study demonstrates that loss of versican from fibroblasts directly impacts pulmonary inflammation by reducing recruitment and accumulation of neutrophils, monocytes, dendritic cells, and mono-macrophages early during IAV infection. Our data support that mesenchymal-derived versican is necessary for neutrophil adhesion to lung fibroblasts as they transit into the lung interstitium and airways from the pulmonary vasculature. While our study provides important new information, it has also raised several important questions that will need to be answered in future investigations. It remains unknown which adhesion molecules on neutrophils, monocytes, and mono-macrophages are responsible for binding these leukocytes to the versican•HA matrix. Additionally, versican-enriched matrices may alter the activation state of leukocytes either directly through interactions between leukocytes and the matrix or indirectly by significantly altering endothelial cell function and activation. Taken together, mesenchymal-derived versican is a key integrator of the early pro-inflammatory host immune response to IAV warranting additional investigation.

**Supplemental Figure S1. UMAP visualization of lung cells and versican expression of mice on days 0-, 3-, and 6-dpi with IAV.**

(A) Uniform manifold approximation and projection (UMAP) plot of the scRNAseq data from uninfected mice. (B) Cell clusters of different colors represent the various populations of cells in the lungs 0, 3, and 6-dpi with IAV. (C) UMAP visualization of Col1a1 and Acta2 normalized counts. (D) UMAP visualization of *Vcan* normalized counts (purple dots) in all cells of the lungs on 0, 3, and 6-dpi with IAV. Grey illustrates cells not expressing *Vcan* mRNA. Abbreviations: AT1, alveolar type 1; AT2, alveolar type 2; AM, alveolar macrophages; LEC, lymphatic endothelial cells; EC, endothelial cells

**Supplemental Figure S2. Col1a2 expression in mesenchymal cells from whole lungs of mice 0-dpi with IAV.** (A) Uniform manifold approximation and projection (UMAP) visualization of mesenchymal cell clusters recovered from whole lungs of mice. (B) Col1a2 normalized counts in the four clusters of mesenchymal cells in control mice (0 dpi) (C) Col1a2 expression in the four clusters of mesenchymal cells in control mice (0 dpi). n =4=5 mice per group.

**Supplemental Figure S3. Histological evidence of tissue injury and alteration of the alveolar-capillary barrier during IAV**

(A) Acute lung injury scores from WT and Col1a2/Vcan^-/-^ mice 9 dpi with IAV. (B) Total protein measured from BAL fluid at 0 (PBS), 3, 6, and 9 dpi with IAV. Values are mean ± SEM, *n* = 3 in (A) and *n* = 4-14 in (B). Mann-Whitney for (A) and two-way ANOVA with Bonferroni’s for multiple comparison test for (B). Abbreviations: BAL, bronchoalveolar lavage

**Supplemental Figure S4. Circulating Leukocytes on 3 dpi with IAV** Total leukocytes, lymphocytes, neutrophils, monocytes, and eosinophils in circulation was determined from whole blood samples collected from Col1a2/Vcan^-/-^ and WT mice 3 dpi with IAV. No differences were observed between Col1a2/Vcan^-/-^ and WT mice using Student’s t-test. Values are mean ± SEM, *n* = 5.

## DATA AVAILABILITY

Data will be made available upon reasonable request.

## ACKNOWLEDGEMENTS

We thank Tom Wight, W. Conrad Liles, and Kristina Adams-Waldorf, for their mentorship and collaboration on this research. In addition, we thank Brian Johnson and the research scientists in the Histology and Imaging Core at UW-South Lake Union for their assistance with measuring inflammatory mediators in the BAL fluid.

## AUTHOR CONTRIBUTION

JEB, WAA, CWF, MYC, FT conceived and designed the research; JEB, MYC, FT, CKC, PW performed experiments; JEB, MYC, FT, CKC, CLM analyzed data; JEB, WAA, CWF, FT, MYC interpreted results of experiments; JEB, CLM prepared figures; JEB drafted manuscript; JEB, MYC, FT, CLM, SRR, CKC, PW, DFB, SAG, PGT, WAA, CWF edited and revised the manuscript; JEB, MYC, FT, CLM, SRR, CKC, PW, DFB, SAG, PGT, WAA, CWF approved final version of the manuscript.

