## Supplemental Figures for "Immunomodulation of the Innate Host Response by Mesenchymal-Derived Versican during Influenza A Virus Infection"

### Supplemental Figure S1.

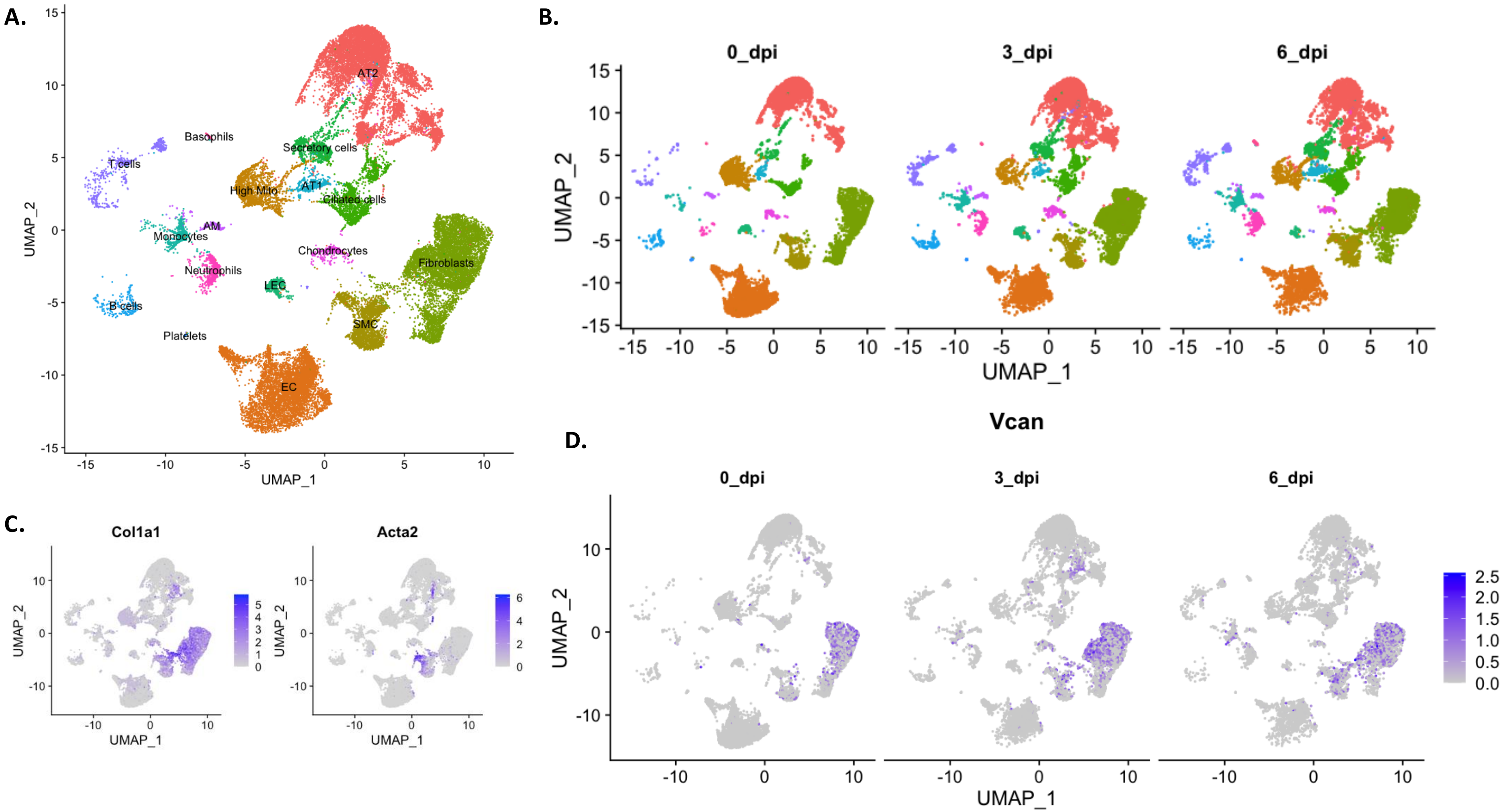

**Supplemental Figure S2.**

A.

B

C.

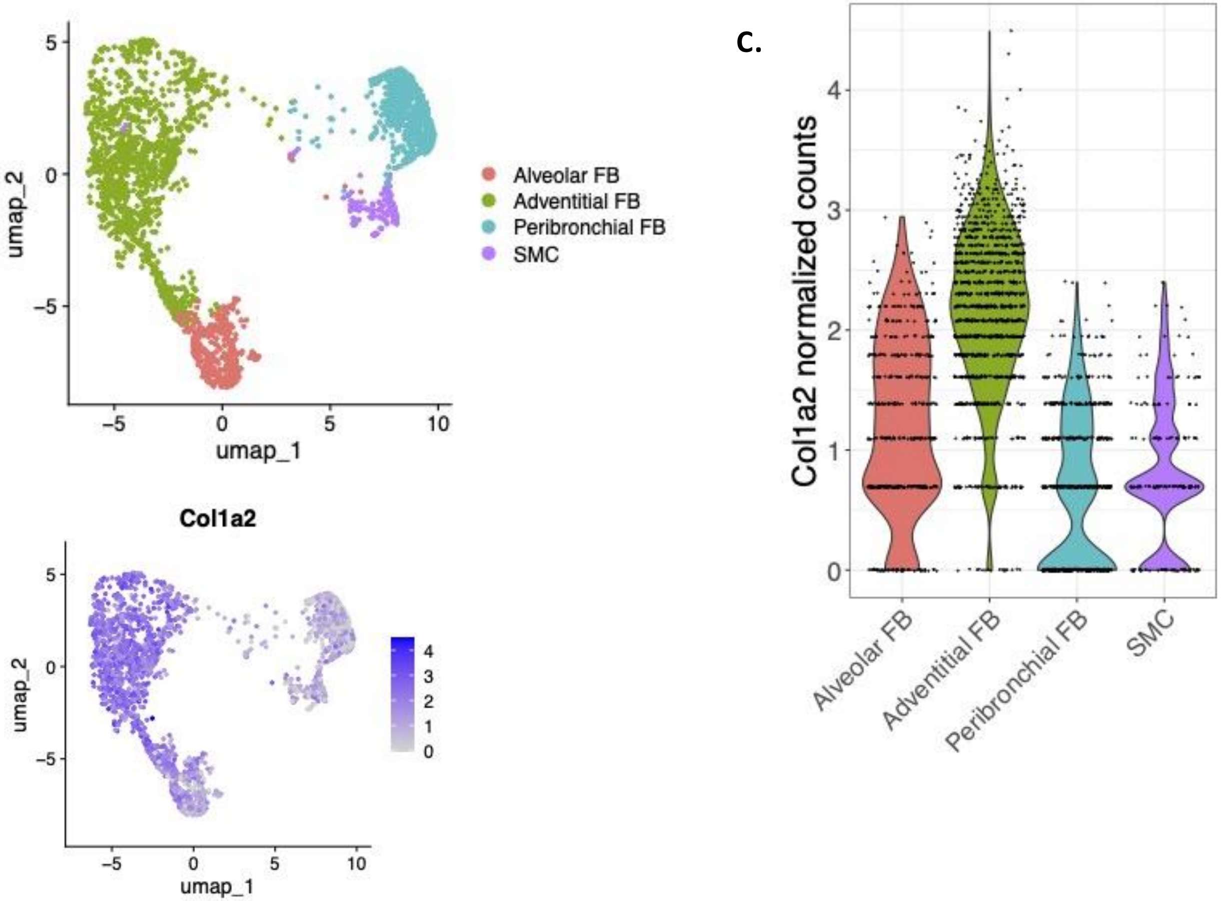

**Supplemental Figure S3.**

**A.**

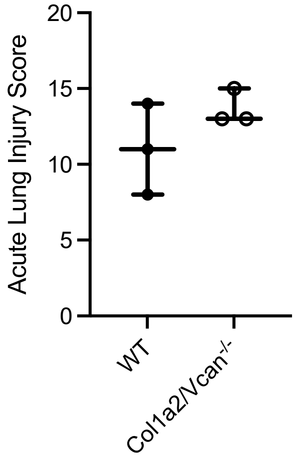


**B.**

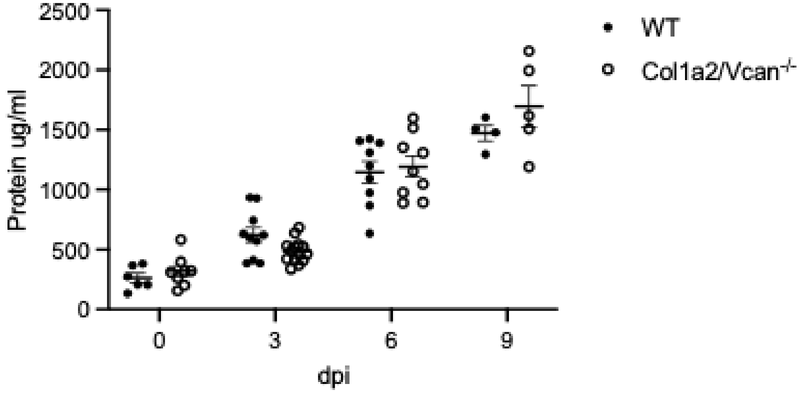

**Supplemental Figure S4.**

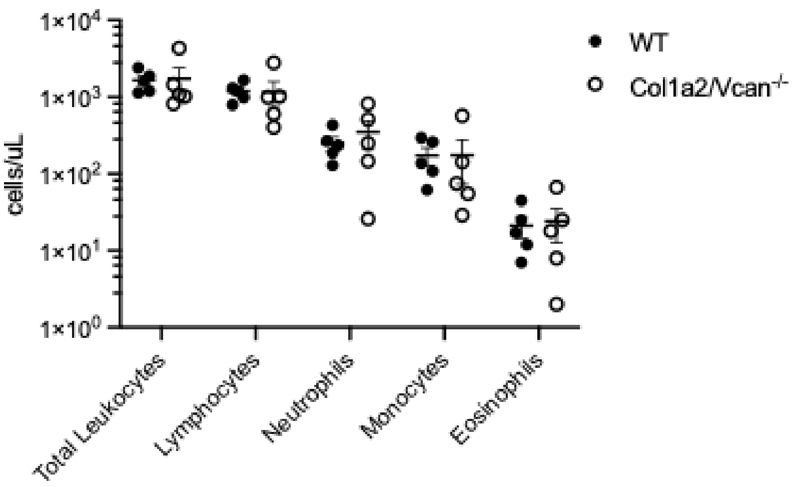
